# Sex-specific co-expression networks and sex-biased gene expression in the salmonid Brook Charr *Salvelinus fontinalis*

**DOI:** 10.1101/305680

**Authors:** Ben J. G. Sutherland, Jenni M. Prokkola, Céline Audet, Louis Bernatchez

## Abstract

Networks of co-expressed genes produce complex phenotypes associated with functional novelty. Sex differences in gene expression levels or in the structure of gene co-expression networks can cause sexual dimorphism and may resolve sexually antagonistic selection. Here we used RNA-sequencing in the paleopolyploid salmonid Brook Charr *Salvelinus fontinalis* to characterize sex-specific co-expression networks in the liver of 47 female and 53 male offspring. In both networks, modules were characterized for functional enrichment, hub gene identification, and associations with 15 growth, reproduction, and stress-related phenotypes. Modules were then evaluated for preservation in the opposite sex, and in the congener Arctic Charr *Salvelinus alpinus*. Overall, more transcripts were assigned to a module in the female network than in the male network, which coincided with higher inter-individual gene expression and phenotype variation in the females. Most modules were preserved between sexes and species, including those involved in conserved cellular processes (e.g. translation, immune pathways). However, two sex-specific male modules were identified, and these may contribute to sexual dimorphism. To compare with the network analysis, differentially expressed transcripts were identified between the sexes, finding a total of 16% of expressed transcripts as sex-biased. For both sexes, there was no overrepresentation of sex-biased genes or sex-specific modules on the putative sex chromosome. Sex-biased transcripts were also not overrepresented in sex-specific modules, and in fact highly male-biased transcripts were enriched in preserved modules. Comparative network analysis and differential expression analyses identified different aspects of sex differences in gene expression, and both provided new insights on the genes underlying sexual dimorphism in the salmonid Brook Charr.

## INTRODUCTION

Understanding how sex bias in gene expression contributes to sexually dimorphic phenotypes, and how this affects fitness is an important area of study to understand genotype-phenotype interactions (Parsch and Ellegren 2013). The development of sex differences in phenotypes can be attributed to differences in the expression of both sex-specific and autosomal genes, caused by hormonal and epigenetic effects (Ellegren and Parsch 2007; Wijchers and Festenstein 2011), heterogamy and imperfect dosage compensation (Parsch and Ellegren 2013), epistatic interactions, and transcriptional network structural differences (Chen *et al.* 2016). Sex-biased gene expression is pervasive in many species and may alleviate sexually antagonistic selection, i.e., the selection for different phenotypes in the different sexes (Wright *et al.* 2018; Rowe *et al.* 2018). Transcriptome regulatory architecture differences between the sexes may contribute to the development of sex-biased gene expression (Chen *et al.* 2016; Wright *et al.* 2018). Sex-biased expression underlies much of phenotypic sexual dimorphism, which can occur without sexually antagonistic selection (e.g. imperfect dosage compensation; Parsch and Ellegren 2013). To connect alleles to phenotypes, typically association studies are applied (Mackay, 2001; Bush and Moore 2012), but these bypass important intermediate regulatory steps such as transcriptome regulation (Mackay *et al.* 2009). To characterize transcriptome regulation, it is important to consider the underlying structure of the network in which genes are co-regulated (Mähler *et al.* 2017).

Constructing gene co-expression networks is often based on correlating transcript abundance across samples (Langfelder and Horvath 2008). A network is comprised of modules, each of which is comprised of a group of genes with correlated expression patterns. Co-expression clustering is a valuable approach to classify and visualize transcriptomic data (Eisen *et al.* 1998). Clustering often groups genes together that have similar cellular functions and regulatory pathways (Eisen *et al.* 1998), although this is not always the case (Gillis and Pavlidis 2012; van Dam *et al.* 2017). Module functions can be predicted based on phenotypic correlations (Filteau *et al.* 2013; Rose *et al.* 2015) or functional enrichment analysis (e.g. Gene Ontology). Genes that are highly connected and central to a module (i.e., hub genes) may be upstream regulators of the module, and potentially more related to the module function (van Dam *et al.* 2017). Hub genes may also be under higher selective constraint than other less connected genes, and as a result may show lower genetic variation and higher phylogenetic conservation (Mähler *et al.* 2017). Network information thus provides novel insight into both gene activity and evolution.

Comparative network analysis across species can advance the understanding of species-specific innovations. For example, comparing brain transcriptome networks between chimpanzee *Pan troglodytes* and human *Homo sapiens* indicates low preservation of modules found in specific brain regions associated with human evolution such as the cerebral cortex (Oldham *et al.* 2006). Comparing networks can highlight potential drivers of phenotypic changes associated with adaptive divergence that can lead to ecological speciation (Filteau *et al.* 2013; Thompson *et al.* 2015). Cross-species network comparisons have also been used to detect gene modules associated with disease (Mueller *et al.* 2017) or with seasonal phenotypic changes (Cheviron and Swanson 2017). These insights are often not possible to obtain through standard gene-by-gene differential expression analysis, which captures a smaller proportion of the variation than differential co-expression analysis (Oldham *et al.* 2006; Gaiteri *et al.* 2014).

Network comparisons can also provide insight on sex differences (Chen *et al.* 2016). Differing structure of networks between the sexes may resolve sexual antagonism through gene regulation (Chen *et al.* 2016). Other genetic architecture solutions to this conflict may include sex-dependent dominance (Barson *et al.* 2015) or maintaining alleles associated with sexual antagonism on sex chromosomes (Blackmon and Brandvain 2017). Comparisons between the sexes are complicated by the fact that sex bias in networks can be tissue-specific, with modules more preserved between sexes in brain and muscle networks than in liver or adipose tissue (van Nas *et al.* 2009; Wong *et al.* 2014). Liver tissue is considered a highly sexually dimorphic tissue, particularly in oviparous species at a reproductive stage (Qiao *et al.* 2016). Although the extent of differences may depend on the tissue of study, network comparisons between sexes can provide new insight into the regulatory underpinnings of sexual dimorphism and antagonism.

Salmonids are an important species of commercial, ecological, and cultural value. This species group is also a model system for studying genome evolution after a whole genome duplication event that occurred approx. 60-88 million years ago (Allendorf and Thorgaard 1984; Crête-Lafrenière *et al.* 2012; Macqueen and Johnston 2014). Charr (*Salvelinus* spp.) are a phenotypically diverse group within Family Salmonidae that are less characterized in terms of transcriptomic and genomic data than other genera (e.g. *Salmo* or *Oncorhynchus*; but see Christensen *et al*. 2018). Brook Charr *Salvelinus fontinalis* is a primarily freshwater species native to Eastern North America. Arctic Charr *S. alpinus* has a circumpolar distribution mainly in the Arctic, and these two lineages diverged approximately 10 million years ago (Horreo 2017). Sexual dimorphism in body size and secondary sexual characteristics is associated with reproductive success in Brook Charr and other salmonids (Quinn and Foote 1994). The largest males are typically dominant and fertilize the majority of a brood (Blanchfield *et al.* 2003). However, smaller sneaker males can also contribute in fertilization, albeit to a lesser extent than the large male. In females however, large body size is more constrained as it is highly associated to fecundity, and smaller life history variants are not expected (reviewed by Fleming 1998). With different optimal reproductive-associated phenotypes between the sexes, this could give rise to sexual dimorphism as well as sexual antagonism.

Here we profile liver transcriptomes of 100 Brook Charr offspring from a single family by RNA-sequencing to characterize co-expression patterns. Transcriptome profiling was conducted shortly (3 h) after the application of an acute handling stressor to all individuals during the reproductive season, increasing variance among individuals. The goals of this study are to i) characterize the sex-specific or preserved modular structure of gene co-expression in liver tissue in Brook Charr; ii) characterize module preservation in the congener Arctic Charr to investigate evolutionary conservation of the networks; iii) connect phenotype and functional category associations to the identified modules; and iv) integrate results from the network analyses with a gene-by-gene differential expression analysis to determine whether the two methods provide different insights on sexual dimorphism in transcriptome architecture.

## METHODS

### Animals and sample collection

Brook Charr used in this study were originally used to construct a low-density genetic map for reproductive (Sauvage, Vagner, Derôme, Audet, and Bernatchez 2012a), growth and stress response QTL analyses (Sauvage, Vagner, Derôme, Audet, and Bernatchez 2012b). They were used to construct a high-density genetic map that was integrated with the other salmonids (Sutherland *et al.* 2016), then used to identify QTL, sex-specific recombination rates, and the Brook Charr sex chromosome (Sutherland *et al.* 2017). The 192 F_2_ individuals were full sibs from a single family resulting from a cross of an F_1_ female and F_1_ male that were from an F_0_ female from a wild anadromous population (Laval River, near Forestville, Québec) and an F_0_ male from a domestic population (Québec aquaculture over 100 years).

F_2_ offspring were raised until 65-80 g and then 21 phenotypes were collected along with several repeat measurements to determine growth rate. Full details on these phenotypes are previously described (Sauvage *et al.* 2012a, 2012b), including sex-specific phenotype averages, standard deviations, and phenotype correlations for all 192 offspring (see Table S1 and Figure S2 from Sutherland et al. 2017). The 15 phenotypes used in the present study to correlate with co-expression modules were maturity, length, weight, growth rate, condition factor, liver weight, post-stress cortisol, osmolality and chloride, change in cortisol, osmolality and chloride between one week before and three hours after an acute handling stress, egg diameter, sperm concentration, and sperm diameter. Fish were anaesthetized with 3-aminobenzoic acid ethyl ester and killed by decapitation as per regulations of Canadian Council of Animal Protection recommendations approved by the University Animal Care Committee, as previously reported (Sauvage *et al.* 2012a). A total of 87% of the 47 females and 96% of the 53 males used for transcriptome profiling were in a reproductive state at the time of the dissection, which was in the fall (smolting occurs in this strain in the spring; Boula *et al.* 2002). Phenotypic sex was determined by gonad inspection (Sauvage *et al.* 2012b). Immediately after decapitation, liver tissue was excised, flash frozen then kept at −80 ^o^C until RNA extraction.

### RNA extraction and library preparation

A total of 100 of the 192 individuals were used for liver transcriptome profiling. Prior to extraction, samples were assigned random order to reduce batch effects on any specific group of samples. Total RNA was extracted from equal sized pieces of liver tissue from approximately the same location on the liver for all samples (0.4 × 0.2 × 0.2 cm; ~1 mg). This piece was rapidly immersed in 1 ml TRIzol (Invitrogen), then placed on dry ice until all samples per batch were prepared (6-12 per extraction round). When all samples were ready, the samples immersed in frozen TRIzol were allowed to slightly thaw for approximately 1 min until beads within the vials were able to move, then the samples were homogenized for 3 min at 20 hz, rotated 180°, and homogenized again for 3 min at 20 hz on a MixerMill (Retsch). The homogenate was centrifuged at 12,000 × g for 10 min at 4 ^o^C. The supernatant was transferred to a new 2 ml tube and incubated for 5 min at room temperature. Chloroform (200 µl) was added to the tube, the tube was shaken vigorously for 15 s and incubated 3 min at room temperature, then centrifuged at 12,000 × g for 15 min at 4 ^o^C. Finally the aqueous layer was carefully transferred to a new centrifuge tube, and into an RNeasy spin column (QIAGEN), as per manufacturer’s instructions with the optional on-column DNase treatment. All samples were quality checked using a BioAnalyzer (Agilent), where all samples had RIN ≥ 8.3 (mean = 9.5), and were quantified using spectrophotometry on a Nanodrop-2000 (Thermo Scientific).

Libraries were prepared using 1 µg of total RNA in the randomized order using TruSeq RNA Sample Prep Kit v2 (Illumina) to generate cDNA as per manufacturer’s instructions using adapters from both Box A and Box B, AMPure XP beads (Agencourt) and a magnetic plate in batches of 8-16 samples per batch. Fragmentation times of 2, 4 and 6 min were tested for optimal size fragmentation and consistency, and as a result of this test, all samples were processed using a 6 min fragmentation time. PCR amplification to enrich cDNA was performed using 15 cycles, as per manufacturer’s instructions. All libraries were quantified using Quant-iT PicoGreen (ThermoFisher) and quality-checked using the BioAnalyzer on High Sensitivity chips (Agilent) for consistent size profiles. Once all samples were confirmed to be high quality and of approximately the same insert size, eight individually tagged samples were pooled in equimolar quantities (80 ng per sample) and sent to McGill Sequencing Center for 100 bp single-end sequencing on a HiSeq2000 (Illumina; total = 13 lanes). Parents (F_1_ individuals) were sequenced in duplicate in two separate lanes.

### RNA-seq mapping

Quality trimming and adapter removal was performed using Trimmomatic (Bolger et al. 2014), removing adapters with the *-illuminaclip* (2:30:10) option and removing low quality reads (< Q2) using *-slidingwindow* (20:2), *-leading* and *-trailing* options. Q2 was used for optimal quantification as previously demonstrated (MacManes 2014). A reference transcriptome for Brook Charr was obtained from the Phylofish database (Pasquier *et al.* 2016). Trimmed reads were mapped against the reference transcriptome with *bowtie2* (Langmead and Salzberg 2012) using *--end-to-end* mode reporting multiple alignments (*-k* 40) for optimal use with eXpress for read count generation (Roberts and Pachter 2013). The multiple alignment file was converted to bam format and sorted by read name using SAMtools (Li *et al.* 2009) and input to eXpress (see full bioinformatics pipeline in *Data Accessibility*).

Read counts were imported into edgeR (Robinson et al. 2010) for normalization and low expression filtering. The smallest library (lib87, 8,373,387 aligned reads) was used to calculate a low-expression threshold. A count-per-million threshold of 10 reads per transcript in this library defined the threshold for transcript retention (cpm > 1.19), as suggested in the edgeR documentation. Any transcript passing this threshold in at least five individuals was retained through the first filtering step for initial data visualization and annotation. Additional low expression filtering was conducted in the differential expression analysis and network analysis (see below). Although transcripts were previously annotated in the Phylofish database (Pasquier *et al.* 2016), each transcript was re-annotated using trinotate (Bryant *et al.* 2017) and *tblastx* against Swissprot (cutoff = 1e^-5^) to obtain as many identifiers as possible for Gene Ontology enrichment analysis. For individual gene descriptions, the re-annotated Swissprot identifier was used primarily, and the Phylofish annotation secondarily.

### Differential expression analysis between sexes

To reduce the number of transcripts with very low expression in the differential expression analysis, we applied a low expression filter of cpm > 1.19 in at least 65% of the individuals from each sex (i.e. ≥ 31 / 47 females or 35 / 53 males). The data was then normalized using the weighted trimmed mean of M-values (TMM Robinson and Oshlack 2010) to generate normalized log_2_ expression levels. Using *model.matrix* from edgeR with sex as the data grouping, a genewise negative binomial generalized linear model (*glmFit*) was fit to each gene. Genes with false discovery rate of ≤ 0.05 and linear fold-change ≥ 1.5 were defined as differentially expressed. These genes were binned into low sex bias (i.e. 1.5 ≤ FC ≤ 4) or high sex bias (i.e. FC > 4) for each sex (negative FC for females, positive for males), as per previous delimitations (Poley *et al.* 2016).

### Weighted gene co-expression network analysis (WGCNA) in Brook Charr

To best estimate associations between modules and phenotypes of interest, sexes were analyzed separately, and then the preservation of each module was evaluated in the opposite sex (Langfelder *et al.* 2011). Due to the independent analysis of each sex, low expression filters (i.e. cpm > 0.5 in at least five individuals) were conducted separately in each sex, then the data normalized as described above. The average library size was 27,896,535 alignments, indicating cpm > 0.5 corresponds to at least 13.95 reads aligning to the transcript for this mean library size. Sex-specific data was used as an input for weighted gene correlation network analysis (WGCNA; Langfelder and Horvath 2008; 2012) in R (R Core Team 2018).

Within each sex, sample outliers were detected and removed by clustering samples based on transcript expression by Euclidean distance and visually inspecting relationships with the *hclust* average agglomeration method of WGCNA (Langfelder and Horvath 2008). Removal of outliers prevents spurious correlations of modules due to outlier values and improves network generation (Langfelder *et al.* 2011). Remaining samples were then correlated with trait data using *plotDendroAndColors*. Network parameters for both female and male networks were defined as per tutorials using unsigned correlation networks (Langfelder and Horvath 2008). Unsigned networks consider the connectivity between identical positive or negative correlations to be equal, and thus genes in the same module may have similar or inverse expression patterns. An optimal soft threshold power (6) was identified by evaluating effects on the scale free topology model fit and mean connectivity by increasing the threshold power by 1 between 1-10 and by 2 between 12-20 (Figure S1), as suggested by Langfelder and Horvath (2008). An unsigned adjacency matrix was generated in WGCNA to identify the 25,000 most connected transcripts to retain for reducing computational load. Then, to further minimize noise and spurious associations, adjacency relationships were transformed to the Topological Overlap Matrix using the *TOMdist* function (Langfelder and Horvath 2008).

Similarity between modules was evaluated using module eigengenes (i.e. the first principal component of the module). Dissimilarity between eigengenes was calculated by signed Pearson correlation as suggested by Langfelder and Horvath (2008) and plotted using *hclust.* When modules were more than 0.75 correlated (dissimilarity 0.25), they were merged as suggested by Langfelder and Horvath (2008). Merged module eigengenes were then tested for associations with phenotypes by Pearson correlation. Notably the sign of the correlation does not necessarily indicate the direction of the relationship between the expression of specific genes in each module and the phenotype because the modules were built using unsigned networks.

Module membership (i.e., the module eigengene-gene correlation) was used to define the top central transcripts for each module (i.e., hub genes). Gene significance (i.e., the absolute value of the trait-gene correlation) was calculated for each transcript against traits weight, specific growth rate, condition factor, hepatosomatic index, change in cortisol, osmolality and chloride from the brief handling stressor, female egg diameter, and male sperm concentration and diameter. Module eigengenes were tested for correlation against traits using Pearson correlation (p ≤ 0.01).

Gene Ontology enrichment analysis of transcripts within each module was conducted using the re-annotated Swissprot identifiers in DAVID Bioinformatics (Huang *et al.* 2009). Heatmaps for modules of interest were generated by using the package *gplots* using the normalized log_2_ cpm data (Warnes *et al.* 2016). Expression values were standardized across samples for each transcript and Pearson correlation was used to cluster transcripts and samples.

To determine sex-specific or sex-conserved modules, module preservation was evaluated by comparing male transcript expression to the generated female modules, and visa-versa, using the *modulePreservation* function of WGCNA. A total of 200 permutations of randomly assigned module labels were used to calculate module preservation rank and Zsummary (Langfelder *et al.* 2011). Low Zsummary scores indicate no preservation (≤ 2), intermediate indicate moderate preservation (2-10) and high scores (≥ 10) indicate strong module preservation (Langfelder *et al.* 2011). The similarity in ranking of modules in terms of preservation in the two different comparative networks (i.e., the opposite sex in Brook Charr and the male Arctic Charr data) was performed by testing the correlation of the module preservation statistic using Spearman rank correlation in R. Module quality was also determined for each module as a measure of module robustness that is characterized by conducting the analysis on multiple random subsets of the original data (Langfelder *et al.* 2011). In addition, cross-tabulation of the proportions of female modules in male modules and visa-versa were performed in R. Cross-tabulation requires similar modular structures of the compared networks, whereas adjacency comparisons directly compare co-expression independent of network topology. All pipelines to analyze the current data are documented and available on GitHub (see *Data Accessibility*).

### Module preservation in Arctic Charr

To compare module preservation between Brook Charr and Arctic Charr *S. alpinus* we used RNA-seq data from 1+ year-old Arctic Charr. The broodstock of this population was reared in hatchery conditions for three generations after being collected from a subarctic, land-locked population in Finland (Lake Kuolimo, 61°16′ N; 27°32′ E). The data were collected from nine male liver samples (fish relatedness not known) from each of 8 ^o^C and 15 ^o^C (total = 18 samples), but due to a large effect of temperature on the transcriptome, and differences between sampling times in the 15 ^o^C group, only the nine samples from the 8 ^o^C, group were used here (normal rearing temperature during summer at the fish hatchery; Figure S2) (Prokkola *et al.* 2018). Fish body mass at 8°C was on average 24.2 g ± standard deviation (S.D.) 10.4 g.

Sample processing was explained fully by Prokkola et al (2018) and briefly described here. In August 2013, fish were euthanized using 200 ppm sodium bicarbonate-buffered tricaine methanesulfonate (MS-222), after which liver samples were collected and flash frozen in liquid nitrogen (Prokkola *et al*. 2018). RNA was extracted from approximately 10 mg of liver tissue using Tri-reagent (Molecular Research Center), and quality checked using a BioAnalyzer (Agilent), with an average identified RNA integrity number of 9.95. Strand-specific cDNA library preparation and sequencing were conducted at Beijing Genomics Institute (BGI Hong Kong) using TruSeq RNA Sample Prep Kit v2 (illumina) and sequenced on an Illumina HiSeq2000 instrument to generate paired-end 100 bp reads. All samples were pooled with unique barcodes across four sequencing lanes. Adapters were removed at BGI, and reads trimmed with Trimmomatic (Bolger et al. 2014) using options *leading* and *trailing* (5) *slidingwindow* (4:15) and *minlen* (36). From samples included in this study, on average 41.7 ± S.D. 7.4 million reads were obtained.

Transcript expression was calculated as above, including using the Brook Charr reference transcriptome for ease of cross-species comparisons. Low expression filtering and normalization for the Arctic Charr data was conducted as above (cpm > 0.5 in at least five individuals). However, a network was not constructed for these samples. Using samples from both temperatures, modules were previously identified (Prokkola *et al.* 2018). Once normalized and input to WGCNA, read counts in Arctic Charr 8°C samples were used to build a gene adjacency matrix, which was then compared against modules generated for female and male Brook Charr samples using the *modulePreservation* function as described above. Caveats regarding this data should be noted, including the smaller sample size, immature state of Arctic Charr, minor differences in rearing environments (albeit both were reared in hatchery conditions), and unknown relatedness.

### Identifying one transcript per gene and assigning chromosome positions

The Brook Charr reference transcriptome (Pasquier *et al.* 2016) possibly contains multiple isoforms for individual genes (Y. Guiguen, *pers. comm.*). Therefore an approach was taken here to reduce multiple genes to a single transcript per gene for enrichment analyses on the sex chromosome or in sex-specific modules. First, the Brook Charr reference transcriptome was aligned to the Atlantic salmon *Salmo salar* chromosome-level genome assembly RefSeq GCF_000233375.1 (Lien *et al.* 2016) using GMAP (Wu and Watanabe 2005). Alignments were converted to an indexed bam retaining only high quality (-q 30) alignments using samtools (Li *et al.* 2009). The indexed bam was converted to a bed file using bedtools *bamtobed* (Quinlan and Hall 2010). Second, the lengths of Brook Charr transcripts were calculated using custom python scripts (see *Data Accessibility*). With the alignment position, lengths, and expression status (expressed or not expressed), a single transcript per contiguous (or overlapping) alignment block on the reference genome was retained using a custom R script (see *Data Accessibility*). For each contiguous alignment block, this script preferentially retained the longest, expressed transcript. All other redundant transcripts, and all that did not align, were not retained. In some cases, a single transcript can align to multiple locations with high mapping quality (MAPQ ≥ 30). Since there was no reason to retain one alignment over another, both alignments were retained in the baseline set for these cases.

The alignment information per retained Brook Charr transcript was used to assign an Atlantic Salmon chromosome identifier to each retained unique transcript. The chromosome information was combined with the module information for all transcripts in each sex-specific network. This analysis was conducted separately for females and males, as expressed genes were in some cases different between the two sexes and therefore so would be the selection of which transcript to retain. The correspondence between the Atlantic Salmon genome assembly accession identifier and the Atlantic Salmon chromosome identifier were obtained from the NCBI genome assembly website (see *Data Accessibility*). In general, correspondence of gene synteny is expected to be similar between Atlantic Salmon and Brook Charr (Sutherland *et al.* 2016). Using the chromosome correspondence table in Sutherland et al. (2016), the Atlantic Salmon chromosome containing the sex chromosome of Brook Charr was identified. For each sex-specific co-expression module, the proportions of Brook Charr genes located on the Atlantic Salmon chromosome that corresponds to the Brook Charr sex chromosome were characterized and compared to the total list of all non-redundant Brook Charr transcripts identified. Fisher exact tests were then used to determine significance for each sex-specific module-sex chromosome combination (i.e., two tests). The chromosome information was also combined with the sex bias fold change values. Finally, the proportion of highly or moderately sex-biased transcripts for females and males were investigated for overrepresentation on the sex chromosome and in sex-specific or conserved modules using Fisher’s exact tests in custom R scripts (see *Data Availability*).

## RESULTS

### Transcriptome overview

Of the total 69,440 transcripts in the Brook Charr reference transcriptome, 51,911 passed initial low expression filters when using all samples together. Low-expression filtering for differential expression analysis (i.e., expressed in at least 65% of individuals from one of two sexes) resulted in the retention of 42,622 transcripts. Low-expression filtering for sex-specific network analysis (i.e., expressed in at least five individuals of the sex of interest) resulted in 50,748 and 50,530 transcripts passing filters in females and males, respectively. When considering each sex individually, most of the expressed genes were expressed in a majority of the samples: females expressed 35,461 transcripts in > 90% of the samples; males expressing 35,714 transcripts in > 90% of the samples (Figure S3).

Hierarchical clustering of samples by gene expression indicated a large effect of sex, where 35 of 47 F_2_ females clustered with the F_1_ female, and 52 of 53 F_2_ males were clustered together with the F_1_ male (Figure 1). As described in the Methods, outliers were removed to avoid spurious network correlations (Langfelder *et al.* 2011), and this included the removal of one male leaving 52 males remaining, and the removal of one group of females that had large liver weight, leaving 35 females remaining (see phenotype liver weight in Figure 1; Figure S4). When these outlier samples were included while constructing the female network, many modules correlated with the liver weight phenotype (*data not shown*), suggesting that these samples were having a large impact on the network.

**Figure 1.**
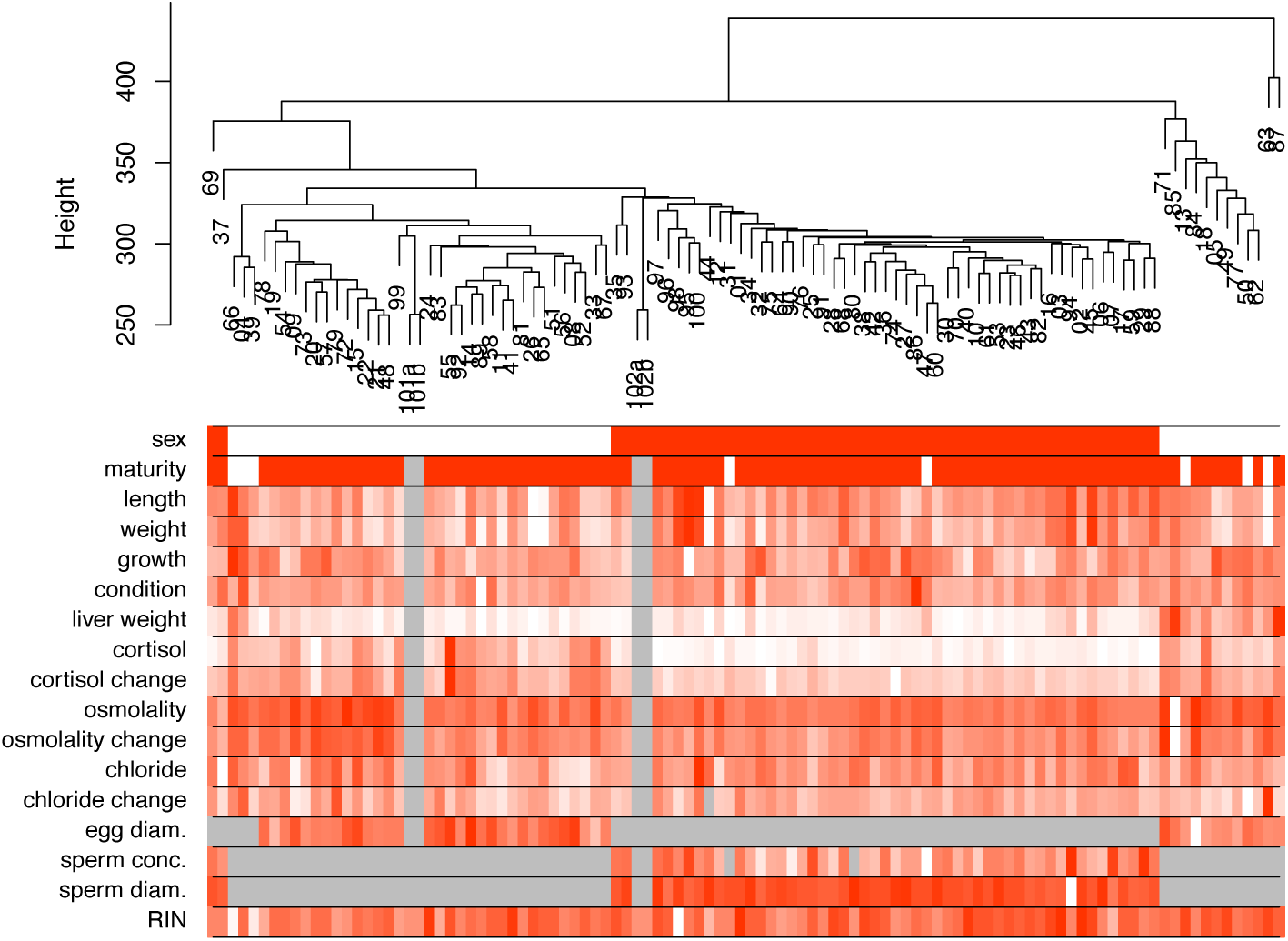
Brook Charr individual samples clustered by gene expression similarity in the liver using all genes with corresponding quantitative trait values shown in the heatmap below the dendrogram with intensity of red reflecting the normalized trait value for that sample. Sex was the largest factor affecting the data (see heatmap sex row; white = females; red = males). Parents were sequenced in duplicate, and clustered with the offspring of their respective sex (see 101ab for mother and 102ab for father; parents have grey missing data values for all phenotypes but sex and RIN). Females with large liver weight clustered outside the other female samples (see on the right-hand side on the liver weight row), and were removed as they were considered outliers (see Methods).

Interestingly, females displayed higher inter-individual variance in gene expression than males, as indicated by the multiple smaller sample clusters of females in the hierarchical clustering relative to the fewer and larger sample clusters of the males (Figure 1). Likewise, six of the phenotypic traits also displayed higher variance in females (significant heteroscedasticity at P < 0.05, Levene’s test): liver weight, cortisol level, osmolality, and change in cortisol, osmolality and chloride levels due to handling (Figure S5). Some of the higher variance in these traits in females was likely explained by maturity, but maturity did not explain all of the variance within the female gene expression, as the different sample clusters observed had a variety of maturity states, and samples that were all determined to be mature were present in different sample clusters (Figure 1). In the following, the sex-specific networks will be presented and compared against the alternate sex and the congener Arctic Charr.

### Network construction and phenotype correlations: female Brook Charr

Highly correlated module eigengenes (r > 0.75) were merged, combining 81 modules into 14 (Figure S6A; Figure S7A). Assigned female modules each contained a range of 77-10,533 transcripts (Table 1). The largest module was *darkred*, with 10,533 transcripts (see Table 1), which included more transcripts than even the unassigned *grey* module (second largest; 5,892 transcripts).

**Table 1.** Female modules shown with the number of transcripts within the module (n), the general category of traits correlated with the module (p ≤ 0.01), the most significantly enriched Gene Ontology category (Biological Process), the Zsummary for preservation of the module in Brook Charr males (BC m) and Arctic Charr males (AC m), as well as the module quality (robustness). Zsummary < 2 is not preserved, 2 < Zsummary < 10 is moderately preserved, and > 10 is preserved. The *grey* module includes unassigned genes and the *gold* module is a random selection of 1000 genes from the assigned modules for testing preservation metrics. Full module-trait correlations are shown in Figure 2, full GO enrichment in Additional File S1, and expanded summaries of this table in Additional File S2.

| Module | n | Traits | GO Enrichment (BP) | Preservation<br>BC m \| AC m |  | Quality |
| --- | --- | --- | --- | --- | --- | --- |
| *green* | 725 | blood | translation | 100 | 24 | 75 |
| *blue2* | 1762 | blood; liver;<br>maturity; RIN | ribonucleoprotein complex<br>biogenesis | 71 | 32 | 58.5 |
| *salmon* | 449 | liver;<br>maturity; RIN | erythrocyte development | 56 | 17 | 42 |
| *darkorange* | 805 | blood | small molecule metabolic<br>process | 48 | 8.8 | 36 |
| *lightsteelblue* | 77 | blood | *none* | 48 | 8.2 | 25.5 |
| *ivory* | 451 | - | ER-associated ubiquitin-<br>dependent protein catabolic<br>process | 37 | 5.4 | 25.5 |
| *thistle2* | 834 | maturity | tissue development | 29 | 3.6 | 36.5 |
| *indianred4* | 1180 | blood;<br>growth; liver;<br>maturity; RIN | membrane assembly | 28 | 2.8 | 22 |
| *coral1* | 689 | liver;<br>maturity; RIN | regulation of cell growth | 25 | 5.2 | 34 |
| *thistle3* | 235 | blood; liver;<br>size | inorganic anion transmembrane<br>transport | 24 | 1.2 | 21 |
| *skyblue1* | 96 | RIN | intracellular signal transduction | 17 | 1.2 | 21 |
| *darkmagenta* | 1072 | blood;<br>maturity | single-organism metabolic<br>process | 13 | 5 | 58 |
| *coral2* | 200 | - | regulation of blood circulation | 11 | 2.8 | 14 |
| *grey* | 5892 | Not a module | Not a module | 9 | 2.4 | -18 |
| *gold* | 1k* | blood; size | Not a module | 4.8 | 0.38 | -0.86 |
| *darkred* | 10533 | blood | protein ubiquitination | 4.5 | -0.55 | 23.5 |

Correlations of module eigengenes with specific phenotypes (n = 15) indicate potential functional associations of the modules (Table 1; Figure 2). The strongest associations of phenotypes to modules were with maturity index, for example with *thistle2* (r > 0.81), and *coral1* (r = −0.83). Although the large liver weight outlier samples were removed prior to network generation, liver weight remained highly correlated with *indianred4* (r = 0.73), *salmon*, *coral* (r = 0.52), and *blue2* (r = −0.64). Growth rate showed similar module correlations to liver weight (Figure 2). Osmolality change was also correlated with *indianred4* (r = 0.56) and *blue2* (r = −0.58), as well as with *darkred*, *thistle3* (r ≥ 0.51), *darkorange*, *green* and *darkmagenta* (r ≥ |-0.49|). Chloride change had no significant associations, but post-stress chloride was correlated with *thistle3* (r = 0.56) and *lightsteelblue* (r = −0.53). Although no modules were significantly associated with cortisol (change or post-stress; p ≥ 0.01), *ivory* was close (r = −0.38; p = 0.03).

**Figure 2.**
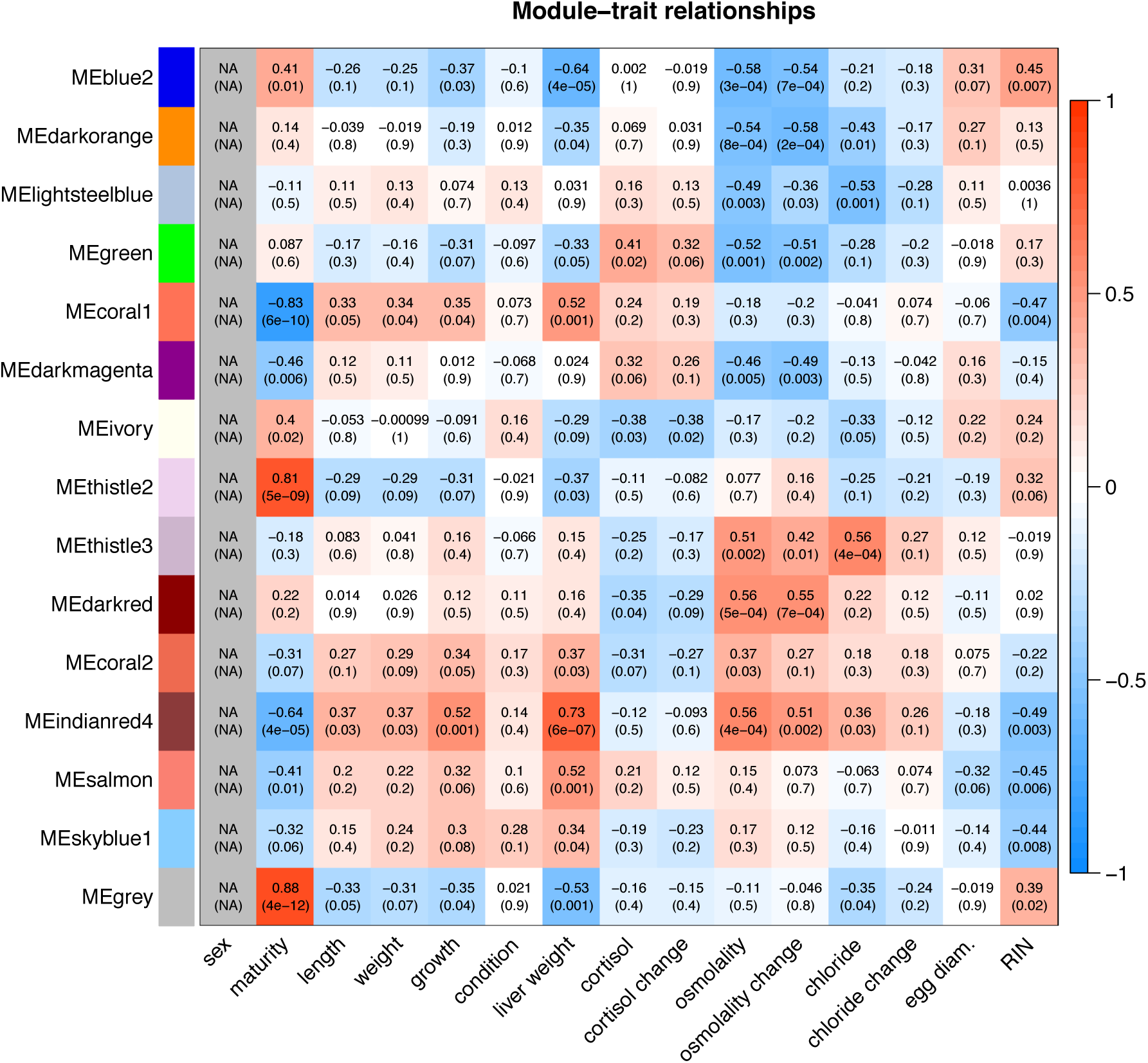
Module-trait relationships for Brook Charr females, estimated with Pearson correlation r-values and p-values. The boldness of color indicates the strength of the relationship. Module-trait correlations are also shown in Table 1 with more general grouping of traits alongside other metrics such as module size and enriched Gene Ontology categories. The male network module-trait relationships are shown in Figure S9.

In order to understand the gene composition of modules, Gene Ontology functional enrichment analysis of the transcripts within each module was conducted (Table 1; Additional File S1). The *salmon* module (correlated with liver weight) was enriched for erythrocyte development. The *blue2* module (liver weight) was associated with ribonucleoprotein complex. *Darkorange*, *green*, and *darkmagenta* modules (all correlated with osmolality change) were enriched for small molecular metabolic process, translation and metabolism functions, respectively. One module did not have significant enrichment of biological processes, *lightsteelblue* (correlated with chloride change).

### Preservation of co-expression: female Brook Charr network

The preservation of co-expression of female Brook Charr modules in Brook Charr males was primarily evaluated using network adjacency comparisons, which are more sensitive and robust than cross-tabulation methods (Langfelder *et al.* 2011). Most female modules were preserved in males and only *darkred* had weak evidence for preservation (Table 1). *Green* was the most conserved (Zsummary = 100), followed by *blue2*, *salmon*, *lightsteelblue* and *darkorange* (Zsummary ≥ 48; Table 1). Modules associated with translation activities were among the highest conserved modules (e.g. *green* and *blue2*).

Published male Arctic Charr liver transcriptome data was then compared to the network to evaluate cross-species module preservation (Prokkola *et al.* 2018). Even with caveats regarding sample size (see Methods), several female Brook Charr modules were highly preserved in Arctic Charr males, including *blue2* (Zsummary = 34), *green* (Zsummary = 24), and *salmon* (Zsummary = 17), also the most conserved in male Brook Charr (Table 1). Other female Brook Charr modules with moderate evidence for preservation in male Arctic Charr included *darkorange* and *lightsteelblue* (Zsummary > 8), which were also highly preserved in male Brook Charr. It is noteworthy that the ranking of preservation of female modules in the Arctic Charr and Brook Charr males is highly similar (Table 1; Spearman rho=0.895; p < 0.00005; Figure S8).

### Network construction, phenotype correlations, and sex-specificity: male Brook Charr

Highly correlated male modules (eigengene correlation r > 0.75) were merged, reducing 44 assigned male modules to 25 (Figure S6B; Figure S7B). Unlike the female network, a large proportion of the male data could not be assigned to a module. The unassigned *grey* module contained 72% of the analyzed transcripts (17,992 transcripts). Assigned modules each contained between 54-1,732 transcripts (Table 2; Additional File S2). Phenotypic correlations with male module eigengenes were tested (Figure S9; see Supplemental Results; summarized in Table 2). Of note were several modules enriched for immunity-related functions, including *darkmagenta* (defense response to virus) and *steelblue* (positive regulation of innate immune response) (Table 2 and Additional File S1). These two immune processes were found to belong to different modules that were not just inversely regulated but rather having somewhat decoupled regulation, given that the network constructed was unsigned and thus the sign of the correlation does not affect whether the genes are grouped (Figure 3A). However, these modules were still correlated even if not grouped into a single module (Figure S7B).

**Table 2.** Male modules with the number of transcripts within the module (n), the general category of traits correlated with the module (p ≤ 0.01), the most significantly enriched Gene Ontology category (Biol. Proc.), the Zsummary for preservation of the module in Brook Charr females (BC f) and Arctic Charr males (AC m), as well as the identified module quality (robustness). Zsummary < 2 is not preserved, 2 < Zsummary < 10 is moderately preserved, and > 10 is preserved. The *grey* module includes unassigned genes and the *gold* module is a random selection of 1000 genes from the assigned modules for testing preservation metrics. Full module-trait correlations are shown in Figure S9, full GO enrichment in Additional File 1, and expanded summaries of this table in Additional File S2.

| Module | n | Traits | GO Enrichment (BP) | Preservation<br>BC f \| AC m |  | Quality |
| --- | --- | --- | --- | --- | --- | --- |
| *yellow* | 339 | blood; liver; size | translation | 58 | 24 | 51 |
| *tan* | 173 | blood; size | *none* | 43 | 18 | 41.5 |
| *brown* | 1732 | blood; sperm | ribonucleoprotein complex biogenesis | 38 | 39 | 72.5 |
| *lightcyan* | 734 | blood; size | organic acid metabolic process | 34 | 21 | 69 |
| *magenta* | 226 | *none* | response to endoplasmic reticulum stress | 32 | 38 | 45 |
| *darkmagenta* | 61 | *none* | defense response to virus | 27 | 16 | 25.5 |
| *darkgreen* | 724 | blood; growth; liver; size | *none* | 24 | 11 | 58 |
| *steelblue* | 65 | blood; growth; sperm | positive regulation of innate immune response | 22 | 6.2 | 26.5 |
| *violet* | 64 | liver | sterol biosynthetic process | 21 | 10 | 29.5 |
| *grey60* | 147 | *none* | secretion by cell | 20 | 11 | 39 |
| *lightsteelblue1* | 55 | *none* | *none* | 15 | 0.85 | 23 |
| *yellowgreen* | 61 | blood | *none* | 13 | 0.24 | 24.5 |
| *turquoise* | 783 | *none* | immune system process | 12 | 14 | 68.5 |
| *orangered4* | 57 | sperm | regulation of apoptotic process | 12 | 2.3 | 26.5 |
| *floralwhite* | 52 | *none* | glutathione metabolic process | 11 | 13 | 25 |
| *paleturquoise* | 64 | blood | response to organonitrogen compound | 10 | 7.8 | 25.5 |
| *lightcyan1* | 791 | blood; sperm | cytokinesis | 9.7 | 11 | 67.5 |
| *sienna3* | 61 | liver; size | extracellular structure organization | 9.6 | 14 | 26.5 |
| *plum1* | 58 | *none* | *none* | 9.6 | 7.4 | 23.5 |
| *mediumpurple3* | 57 | size | ribonucleoprotein complex assembly | 8.4 | 5.3 | 20.5 |
| *darkolivegreen* | 138 | blood | *none* | 7.6 | 11 | 28.5 |
| *skyblue* | 78 | RIN | regulation of transcription, DNA-templated | 6.7 | 5.8 | 28 |
| *grey* | 17992 | *Not a module* | *Not a module* | 5.5 | 2.3 | -14 |
| *gold* | 1k* | *Not a module* | *Not a module* | 4.9 | 3.2 | -0.035 |
| *green* | 334 | size | translation | 4.5 | 3.1 | 39 |
| *darkgrey* | 100 | size | small molecule metabolic process | 1.8 | -0.28 | 31.5 |
| *ivory* | 54 | blood | neurogenesis | 0.81 | 0.22 | 24 |

**Figure 3.**
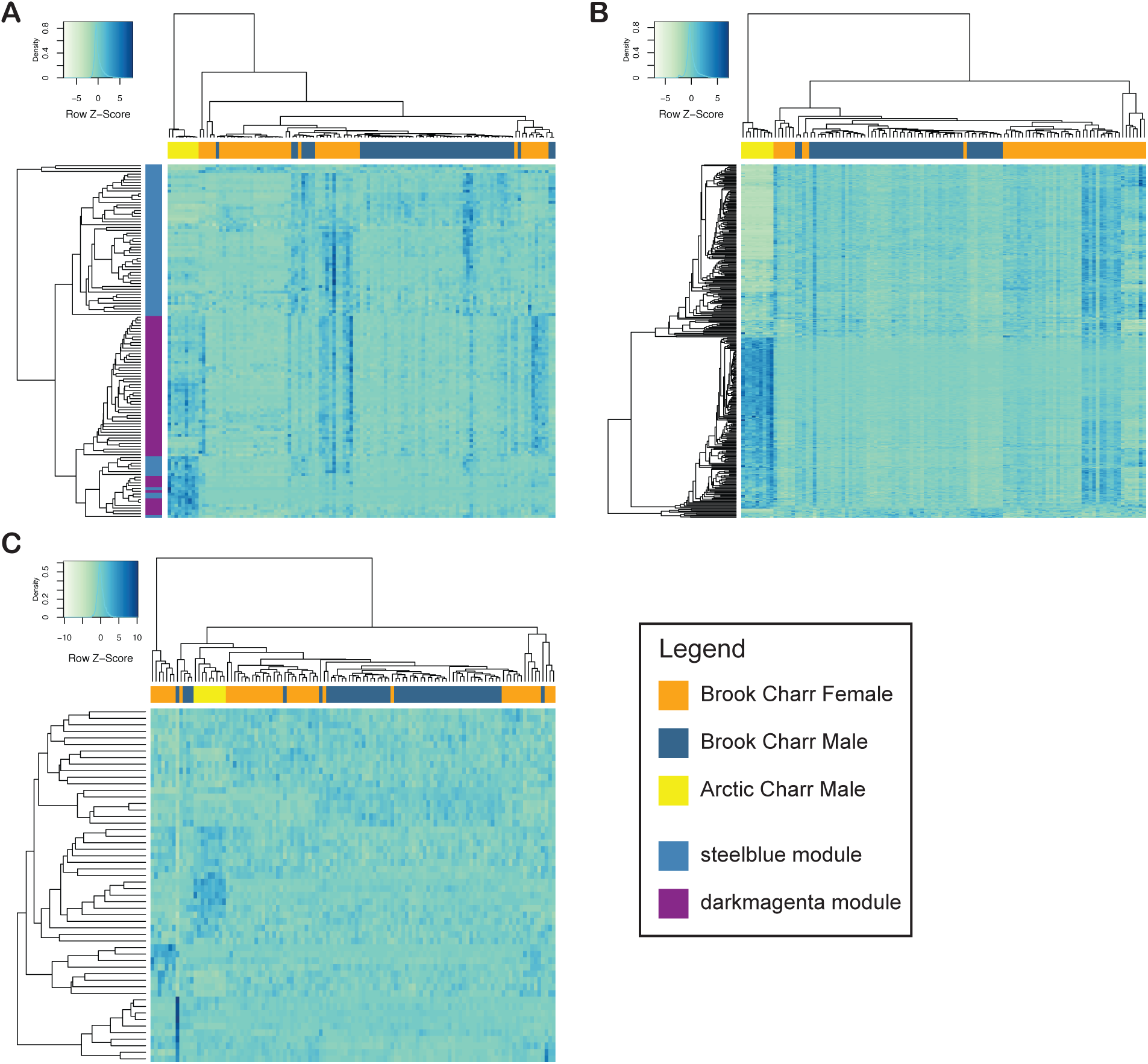
Heatmaps of normalized transcript expression values, clustering both samples and transcripts using Pearson correlation within (A) two immunity-related male modules, *steelblue* and *darkmagenta*, which are related to innate immunity and innate antiviral immunity, respectively, (B) the preserved male module *yellow* (translation), and (C) the male-specific *ivory* (transcription factor activity). Samples are shown on the horizontal with colors corresponding to the three categories of samples in the legend. The two modules shown in (A) are colored on the vertical based on the cluster in which the transcript is contained, as shown in the legend.

Preservation of male modules in females was also evaluated, but in contrast to what was observed in the preservation of female modules in males (see above), many of the male modules were weakly to moderately preserved in the females. Highly preserved modules included *yellow* (Zsummary = 58; Figure 3B) and *brown* (Zsummary = 38), *tan* (Zsummary = 43), and *lightcyan* (Zsummary = 34). Some modules were less preserved, and therefore more sex-specific, than even the randomly generated *gold* module (Table 2) and the unassigned *grey* module, including *green* (translation and size; Zsummary = 4.5), *darkgrey* (mitochondrial membrane; Zsummary = 1.8) and *ivory* (transcription factor activity; Zsummary = 0.8; Figure 3C). Preserved modules *yellow* and *brown* were enriched for ribosomal or translation-related functions, as was the more sex-specific *green* module (Table 2). *Ivory*, the most sex-and species-specific male module (see below; Table 2; Figure 3C) was enriched for neurogenesis in GO biological process, but also transcription factor activity in GO molecular function (Additional File S1). Other non-preserved or lowly preserved modules were enriched for membrane activity including *darkgrey* (mitochondrial inner membrane) and *lightcyan1* (membrane organization; Additional File S1). Preservation of male Brook Charr modules was also explored in Arctic Charr males. Similar to that observed in the female modules, when a male Brook Charr module was preserved in female Brook Charr, it was also often preserved in male Arctic Charr (Table 2; Spearman rho=0.69; p < 0.0005; Figure S8).

### Sex-biased transcripts, sex-specific modules, and the sex chromosome

To further understand the relation between sex-bias in gene expression and sex-specificity in network architecture, a gene-by-gene differential expression analysis between the sexes was conducted. Of the 42,622 expressed transcripts, 6,983 (16.4%) were differentially expressed (FC ≥ 1.5; glmFit FDR ≤ 0.05). Female-biased genes included 3,989 moderately (1.5-4-fold) and 236 highly biased (>4-fold) transcripts. Highly biased transcripts included known sex-biased genes such as *vitellogenin*, and *zona pellucida sperm-binding proteins*. Male-biased genes included 2,638 moderately and 120 highly biased transcripts, including *semaphorin-3F* (most highly male-biased transcript). For a complete list, see Additional File S3.

Interestingly, sex-biased transcripts were not overrepresented in female or male sex-specific modules (Table 3). However, unexpectedly, highly male-biased genes were overrepresented in highly preserved modules (36 transcripts in highly preserved modules, 97% of the highly sex-biased transcripts) in comparison to the overall percentage in the network (4,886 transcripts in highly preserved modules, 76.5% of network transcripts; Table 3; two-sided Fisher’s exact test p = 0.0013). The female data was more similar between the highly sex-biased transcripts and the entire network (45.2% and 46.6%, respectively). Sex-specific modules were not enriched on the sex chromosome (Fisher’s exact test p > 0.5), including male modules *ivory* and *darkgrey* (Table S1).

**Table 3.**
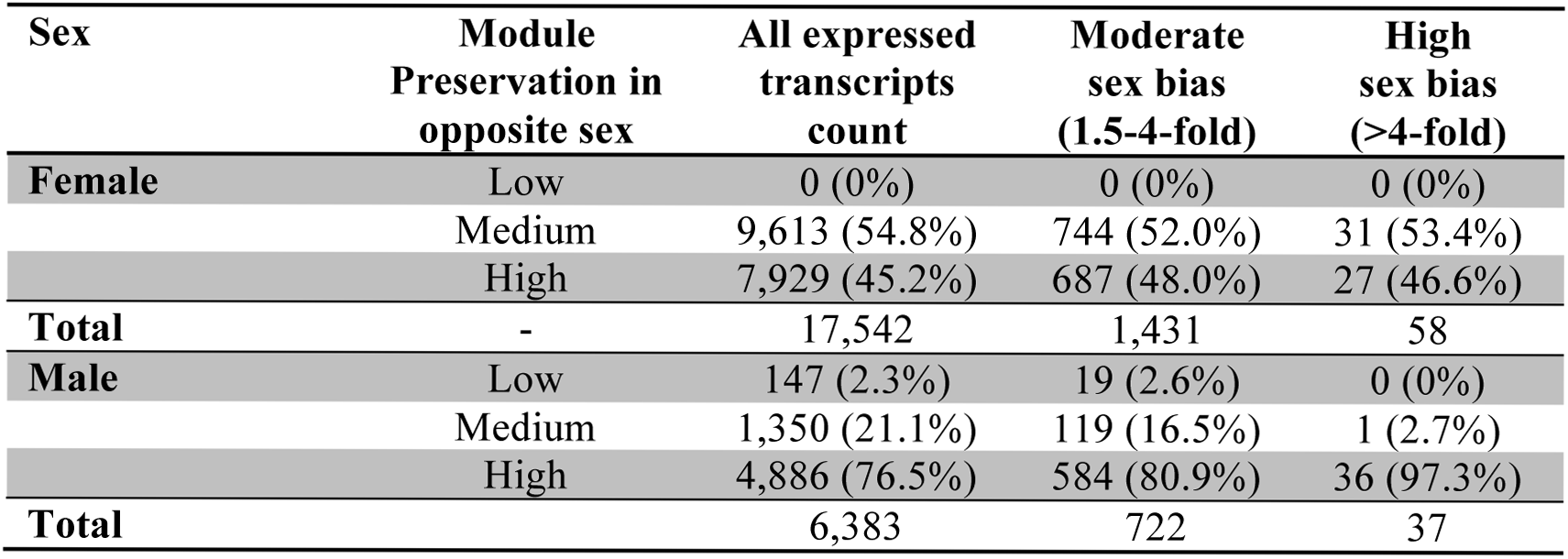
Sex-biased transcript presence in modules that are either unique to each sex (low module preservation), or moderately or highly preserved, along with the number and percentage of the transcripts within each sex’s sex-biased transcript category (e.g. female high sex-bias). These counts only include expressed transcripts that are assigned to modules in the network analysis for each sex.

| Sex | Module Preservation in opposite sex | All expressed transcripts count | Moderate sex bias (1.5-4-fold) | High sex bias (>4-fold) |
| --- | --- | --- | --- | --- |
| Female | Low | 0 (0%) | 0 (0%) | 0 (0%) |
|  | Medium | 9,613 (54.8%) | 744 (52.0%) | 31 (53.4%) |
|  | High | 7,929 (45.2%) | 687 (48.0%) | 27 (46.6%) |
| Total | - | 17,542 | 1,431 | 58 |
| Male | Low | 147 (2.3%) | 19 (2.6%) | 0 (0%) |
|  | Medium | 1,350 (21.1%) | 119 (16.5%) | 1 (2.7%) |
|  | High | 4,886 (76.5%) | 584 (80.9%) | 36 (97.3%) |
| Total |  | 6,383 | 722 | 37 |

Of the 47 non-overlapping, highly male-biased transcripts assigned to chromosomes, five were on the sex chromosome (10.6%), relative to 779 on the sex chromosome of the 12,934 expressed in males (Ssa09; 6.0%). However, this difference was not significant (one-tailed Fisher’s exact test p = 0.15). Furthermore, moderately male-biased transcripts were not enriched on the sex chromosome (5.3%) relative to all expressed transcripts (6%), nor were highly or moderately female-biased transcripts (high bias = 4.1%; moderate bias = 5.8%).

## DISCUSSION

Gene co-expression produces complex phenotypes and may underlie key aspects of phenotypic evolution (Filteau *et al.* 2013) and sexual dimorphism (Chen *et al.* 2016). One of the main challenges in producing females and males is developing different phenotypes from largely the same set of genes (Rowe *et al.* 2018). Transcriptome architecture may provide a solution to this challenge. In this study, co-expression networks for both female and male Brook Charr liver transcriptomes were characterized and compared to each other. Although the female network had much higher module assignment of expressed transcripts, in general most modules were preserved between the sexes. Two sex-specific modules were identified in males that may provide insight on the evolution of gene expression and phenotypic sexual dimorphism. Sex bias was observed in 16% of the expressed transcripts, and surprisingly these sex-biased transcripts were not overrepresented in sex-specific modules. This indicates the value of using these two different approaches given that different information was obtained from each.

### Sex differences in co-expression networks and cross-species preservation

Females and males clustered distinctly in unsupervised clustering by gene expression, as has been observed in other studies of fish liver at a reproductive stage (Qiao *et al.* 2016). Interestingly, there was a large difference in the number of transcripts assigned to modules between the sexes; of the 25 k most connected transcripts, females had 76% transcripts assigning to a module, and males only 28%. Importantly, this observation coincides with higher inter-individual variation in gene expression in females than in males (Figure 1) and higher inter-individual variation in six phenotypes in females than in males (Figure S5). When inter-individual variation in gene expression is low, transcripts will not be as well clustered, because without variance there can be no co-variance for clustering algorithms to act upon (Tritchler *et al.* 2009). Therefore, the lower variance in male gene expression may have resulted in the observed lower module assignment. The observation of lower variation in phenotypes is an interesting result that highlights similarities between transcriptomic and phenotypic expression. Building networks in both sexes allowed for the identification of this lower assignment to modules in males, which may have not been noted otherwise, as female modules generally were scored as preserved in males. Furthermore, this allowed the identification of many male network modules with seemingly important and different functional associations that in the female network were all grouped together into one very large module (i.e., *darkred*). The reason for the grouping of these multiple male modules all together into a single female module is not clear, but could be due to an increased effect of maturity on the data in females that may be swamping out the other more subtle covariances in the data. This further indicates the value of doing separate analyses in each sex, in order to avoid signal being overwhelmed by phenotypes that impact the data more in one sex than the other. It will be valuable to inspect sex-specific module generation in other salmonids and in tissues other than liver to understand the generality of these sex differences and associations to inter-individual variance in gene expression and phenotypes.

Our observations confirm previous findings that co-expression patterns are often preserved between sexes or closely related species (van Nas *et al.* 2009; Wong *et al.* 2014; Cheviron and Swanson 2017). Here, highly preserved modules between the sexes were often comprised of genes within pathways involved in conserved functions. The most preserved modules between the sexes and species were involved in basic cellular processes and included many co-expressed subunits of a multiple subunit protein complex, such as translation machinery. Multiple subunits and functionally related genes have long been known to cluster together by co-expression (Eisen *et al.* 1998). Immunity-related modules were also preserved between the sexes, with co-expression patterns similar to those observed previously in salmonids (Sutherland, Koczka, *et al.* 2014). Considering the importance of immune function to both sexes, it is not surprising that immunity modules are preserved between sexes.

The male-specific module *darkgrey* and the lowly preserved *green* module were both associated with size, which can be sexually dimorphic in salmonids and is associated to breeding success in males (Blanchfield *et al.* 2003). In comparison, no female modules were associated to length and weight, further suggesting that these more male-specific modules could have a role in producing a sexually dimorphic phenotype. The other male-specific module, *ivory*, may contribute to sexual dimorphism and resolution of sexual antagonism by being an upstream controller of different programs, as it is enriched for transcription factor activity and hub genes as putative transcription factors. Hub genes of *ivory* include genes from the *wnt* protein family. Wnt signaling is associated with gonad differentiation and shows sex-specific expression in several studies in mammals and fish (Vainio *et al.* 1999; Nicol and Guiguen 2011; Sreenivasan *et al.* 2014; Böhne *et al.* 2014). Future studies could investigate whether transcription factors from the sex-specific *ivory* module control expression of transcripts found to be sex-biased here once transcription factor binding sites are characterized in this species.

The presence of sex-specific modules and sexually dimorphic gene expression in the liver corresponds with what is known about sex hormones produced in the gonads, as these hormones have been shown to regulate a significant proportion of the liver transcriptome in mouse (van Nas *et al.* 2009). In a large-scale transcriptome study in humans, the liver was not one of the most sexually dimorphic in terms of sex-biased genes (Chen *et al.* 2016). In oviparous species at a reproductive stage however, this tissue is highly sexually dimorphic, given the role in females for producing oocyte constituents (e.g. *vitellogenins*, *zona pellucida* proteins) (Qiao *et al.* 2016). Some of the strongest phenotypic associations of female modules were to maturity. These associations may reflect the effects of sex hormones such as estradiol, which controls reproduction and has a strong influence on transcription in fish (Garcia-Reyero *et al.* 2018). There were no male modules associated to maturity, but there were only six females and two males retained in the analysis that were immature, which prevents a comparison of maturation-related transcripts between the sexes.

The ranking of module preservation levels in both the opposite sex and in Arctic Charr was often similar, suggesting evolutionary conservation for many gene co-expression modules. Even with the lower sample size in Arctic Charr, moderate and high preservation was identified for eight and three of the female modules (n = 14), respectively, and for seven and 14 of the male modules (n = 25), respectively. Modules preserved across species with significant phenotypic correlations may be worthwhile to investigate further regarding their contribution to phenotypes such as growth rate, reproduction, stress response, and immunity. For example, the preserved module in the male network, *turquoise*, was enriched for immunity and marginally associated with growth (p = 0.05), phenotypes known to trade-off (Lochmiller and Deerenberg 2000; van der Most *et al.* 2010).

Many sex-biased transcripts were identified (n = 6,983 transcripts), but only 154 transcripts were found in sex-specific modules identified through the network comparison approach. Sex-biased expression and sex-specific networks are not always overlapping phenomena (Chen *et al.* 2016). This highlights the large differences between these approaches, but in general they together provided a more comprehensive result than either in isolation. The sex bias analysis found that neither male-biased nor female biased genes were significantly overrepresented on the sex chromosome, but power to detect this may have been reduced by the use of a reference genome of a related species rather than the target species. In other species, male-biased transcripts are more often associated with migration to the sex chromosome (Rowe *et al.* 2018), and although there was a trend towards this for the highly male-biased transcripts here, it was not significant. The relationship of sex chromosomes, sex-biased gene expression and sexual dimorphism is not yet well established (Dean and Mank 2014), and this study is an example of integrating these multiple aspects for improved understanding of the role of transcriptomics in generating sex differences.

### Case study: modules separated by immune response type

To demonstrate the utility of this network approach in investigating specific phenotypes, the following is an analysis of modules associated with immunity in the male network. Separate modules were identified for immune functions involving innate antiviral genes (i.e., male *darkmagenta*) and innate immunity C-type lectins (i.e., male *steelblue*). This is of large interest considering that these types of immune responses have been observed to respond inversely, where pathogen recognition receptors (e.g. C-type lectins) are up-regulated and innate antiviral genes down-regulated in the anterior kidney during ectoparasite infection (Sutherland, Koczka, *et al.* 2014) and pathogen recognition receptors are up-regulated and innate antiviral genes are down-regulated in gill tissue during out-migration of steelhead trout *Oncorhynchus mykiss* smolts (Sutherland, Hanson, *et al.* 2014). Even if the genes are not the same between these studies and ours (i.e., no 1:1 association of orthologs has been done for these datasets), the observation of similar functions in two different modules in the present study may indicate that these functions are hardwired into different modules given that no known infection is occurring within these samples. It is important to note that here unsigned networks were used, and therefore if the two immune response types were completely inversely regulated, they would belong to the same module, which was not observed here. These two modules may therefore not be completely under the same regulatory control as they are not completely inversely correlated. This is a new observation in the regulation of these different immune system processes in salmonids. This is an important avenue for further study given the relevance of these genes to immune responses against pathogens, and the potential response outcomes of co-infection occurring between parasitic and viral agents in nature.

The immune response modules observed here (i.e., male *darkmagenta*, *steelblue*, and *turquoise*) were all considered as highly preserved between the sexes and moderately to highly conserved in Arctic Charr. It will be valuable to see if these three modules or the genes within them have conserved expression patterns in other species as these may have important roles in defense responses. The tissue was in a post-stress state, which could affect the induction of immune responses, and so additional observations, such as in a resting state, will be valuable. It is possible that the co-expression viewed in these (and other) modules comes from the occurrence of a specific cell type that is present in different levels in the sampled tissue in different individuals. Single-cell RNA-sequencing of immune cells, or *in situ* gene expression hybridization techniques could address some of these questions. Further, to better understand the immunity related modules, it may also be valuable to use a microbe profiling platform alongside transcriptome studies of wild sampled individuals to best understand co-infection details (e.g. Miller *et al.* 2016). Nonetheless, the characterization of salmonid co-expression modules will be strengthened when additional analyses are conducted with a broader range of species, once orthologs are identified among the species.

### Future comparative approaches and salmonid transcriptome network evolution

When similar datasets are produced in other salmonids, it will be valuable to identify whether the preserved modules are conserved outside of the genus *Salvelinus*. However, importantly this will require identification of 1:1 orthologs among the species, which would enable cross-species analyses. Salmonid ortholog identification across reference transcriptomes has recently been conducted for Atlantic Salmon, Brown Trout *Salmo trutta*, Arctic Charr, and European Whitefish *Coregonus lavaretus* (Carruthers *et al.* 2018), as well as Northern Pike (*Esox lucius*), Chinook Salmon (*O. tshawytscha*), Coho Salmon (*O. kisutch*), Rainbow Trout (*O. mykiss*), Atlantic Salmon, and Arctic Charr (Christensen *et al.* 2018). This type of approach, combined with non-redundant reference transcriptomes will be invaluable in future studies to enable cross species comparisons.

If modules are indeed largely conserved between species, as our study suggests within *Salvelinus* liver, this would indicate that large-scale rewiring of baseline transcription networks has not occurred since the base of the lineage. Species-specific modules will be highly valuable to investigate to better understand transcriptional architecture underlying phenotypic differences between the species. The largest amount of rediploidization is thought to have occurred in the salmonids at the base of the lineage (Kodama *et al.* 2014; Lien *et al.* 2016), although a substantial proportion of ohnologs experienced lineage-specific rediploidization post-speciation events later in evolutionary time (Robertson *et al.* 2017). Given the large potential impact that divergence in regulatory regions or epigenetic signatures can have on gene expression, one could expect large lineage-specific changes in co-expression networks. The impact of the genome duplication and rediploidization on transcriptome networks, including lineage specific changes are important avenues for future study.

## CONCLUSIONS

Co-expression networks and sex-biased expression of female and male Brook Charr liver in a reproductive season shortly after an acute handling stressor were characterized in the present study. Results support previous observations of moderate to high preservation of modules between sexes and closely related species. Highly preserved modules were involved in basic cellular functions and immune functions. Sex-specific modules identified only in the male network were enriched for transcription factor activities and associated with sex-biased or potentially sexually antagonistic phenotypes, such as body size. Higher assignment of transcripts to modules was identified in the female network, potentially due to higher inter-individual variance in gene expression and phenotypes. Important physiological functions such as immunity response types were captured by this analysis, identifying not only inverse regulation between two immunity responses but potentially decoupled regulation, which has implications for responses to co-infections and requires further study. This dataset has provided new insights into the transcriptome network structure differences between sexes and has pointed towards individual genes and gene modules that may be involved with generating sexually dimorphic phenotypes and potentially alleviating sexually antagonistic selection.

## Supporting information

Supplemental Results

## DATA ACCESSIBILITY

Brook Charr Phylofish transcriptome assembly (Pasquier *et al.* 2016): http://phylofish.sigenae.org/ngspipelines/#!/NGSpipelines/Salvelinus%20fontinalis

Atlantic Salmon genome assembly (Lien *et al.* 2016): https://www.ncbi.nlm.nih.gov/assembly/GCF_000233375.1

Complete and documented bioinformatics and analysis pipeline: https://github.com/bensutherland/sfon_wgcna

## SUPPLEMENTAL INFORMATION

**Supplemental Results.** Additional results including figures and tables that support the main text.

**Additional File S1.** All enrichment analyses for Gene Ontology biological process and molecular function categories. Includes all enrichment results with five or more genes in the category and with an enrichment p ≤ 0.01 (no multiple test correction), and Swissprot identifiers within the category.

**Additional File S2.** Overview table for female and male modules, including module size, trait association, GO enrichment, preservation Zsummary and medianRank, and cross-tabulation results.

**Additional File S3.** Differential expression results for the sex-bias analysis, including log_2_ fold change values, p-values, identifiers and gene descriptions when available.

**Additional File S4.** Transcripts and module assignment in the female network along with Gene Significance and Module Membership values.

**Additional File S5.** Transcripts and module assignment in the male network along with Gene Significance and Module Membership values.

**Additional File S6.** Estimated counts file from eXpress used as input for network and differential expression analyses before low expression filtering and normalization.

**Additional File S7.** Interpretation file providing all phenotypes for each sample, used in the analysis pipeline.

## ACKNOWLEDGEMENTS

Thanks to Guillaume Côté for laboratory assistance, Jérémy Le Luyer and Marie Filteau for discussions about WGCNA, Eric Normandeau for discussion on gene annotation, Yann Guiguen for discussion on details of transcriptome assembly, Phineas Hamilton for statistical discussion, and Aimee-Lee Houde, three anonymous reviewers and the Editors for comments on the manuscript. This work was funded by a Fonds de Recherche du Québec (FRQ) Nature et Technologies research grant awarded to Céline Audet, Louis Bernatchez, and Nadia Aubin-Horth; a grant from the Société de Recherche et de Développement en Aquaculture Continentale (SORDAC) awarded to L.B. and C.A. J.M.P. is supported by the Finnish Cultural Foundation. During this work, B.J.G.S. was supported first by a Natural Sciences and Engineering Research Council of Canada postdoctoral fellowship and subsequently by an FRQ Santé postdoctoral fellowship.

## REFERENCES

Allendorf, F. W., and G. H. Thorgaard, 1984 Tetraploidy and the evolution of salmonid fishes, pp. 1–53 in Evolutionary genetics of fishes, edited by B. J. Turner. Plenum Publishing Corporation, New York.

Barson, N. J., T. Aykanat, K. Hindar, M. Baranski, G. H. Bolstad et al., 2015 Sex-dependent dominance at a single locus maintains variation in age at maturity in salmon. Nature 528: 405–408.

Blackmon, H., and Y. Brandvain, 2017 Long-term fragility of Y chromosomes is dominated by short-term resolution of sexual antagonism. Genetics genetics.300382.2017–9.

Blanchfield, P. J., M. S. Ridgway, and C. C. Wilson, 2003 Breeding success of male brook trout (Salvelinus fontinalis) in the wild. Mol. Ecol. 12: 2417–2428.

Bolger, A. M., M. Lohse, and B. Usadel, 2014 Trimmomatic: a flexible trimmer for Illumina sequence data. Bioinformatics 30: 2114–2120.

Boula, D., V. Castric, L. Bernatchez, and C. Audet, 2002 Physiological, endocrine, and genetic bases of anadromy in the Brook Charr, Salvelinus fontinalis, of the Laval River (Québec, Canada). Environ Biol Fish 22: 229–242.

Böhne, A., T. Sengstag, and W. Salzburger, 2014 Comparative transcriptomics in East African Cichlids reveals sex-and species-specific expression and new candidates for sex differentiation in fishes. Genome Biology and Evolution 6: 2567–2585.

Bryant, D. M., K. Johnson, T. DiTommaso, T. Tickle, M. B. Couger et al., 2017 A tissue-mapped Axolotl de novo transcriptome enables identification of limb regeneration factors. Cell Reports 18: 762–776.

Bush, W. S., and J. H. Moore, 2012 Chapter 11: Genome-Wide Association Studies. PLoS Comput Biol 8: e1002822.

Carruthers, M., A. A. Yurchenko, J. J. Augley, C. E. Adams, P. Herzyk et al., 2018 De novo transcriptome assembly, annotation and comparison of four ecological and evolutionary model salmonid fish species. BMC Genomics 19: 32.

Chen, C.-Y., C. M. Lopes-Ramos, M. L. Kuijjer, J. N. Paulson, A. R. Sonawane et al., 2016 Sexual dimorphism in gene expression and regulatory networks across human tissues. bioRxiv 1–34.

Cheviron, Z. A., and D. L. Swanson, 2017 Comparative transcriptomics of seasonal phenotypic flexibility in two North American songbirds. Integr. Comp. Biol. 57: 1040–1054.

Christensen K. A., Rondeau E. B., Minkley D. R., Leong J. S., Nugent C. M., Danzmann R. G., Ferguson M. M., Stadnik A., Devlin R. H., Muzzerall R., Edwards M., Davidson W. S., Koop B. F., 2018 The Arctic charr (Salvelinus alpinus) genome and transcriptome assembly. PLoS ONE 13: e0204076–30.

Crête-Lafrenière, A., L. K. Weir, and L. Bernatchez, 2012 Framing the Salmonidae family phylogenetic portrait: a more complete picture from increased taxon sampling. PLoS ONE 7: e46662.

Dean, R., and J. E. Mank, 2014 The role of sex chromosomes in sexual dimorphism: discordance between molecular and phenotypic data. Journal of Evolutionary Biology 27: 1443–1453.

Eisen, M. B., P. T. Spellman, P. O. Brown, and D. Botstein, 1998 Cluster analysis and display of genome-wide expression patterns. Proceedings of the National Academy of Sciences 95: 14863–14868.

Ellegren, H., and J. Parsch, 2007 The evolution of sex-biased genes and sex-biased gene expression. Nat Rev Genet 8: 689–698.

Filteau, M., S. A. Pavey, J. St-Cyr, and L. Bernatchez, 2013 Gene coexpression networks reveal key drivers of phenotypic divergence in Lake Whitefish. Molecular Biology and Evolution 30: 1384–1396.

Fleming, I. A., 1998 Pattern and variability in the breeding system of Atlantic salmon (Salmo salar), with comparisons to other salmonids. Can. J. Fish. Aquat. Sci. 55: 59–76.

Gaiteri, C., Y. Ding, B. French, G. C. Tseng, and E. Sibille, 2014 Beyond modules and hubs: the potential of gene coexpression networks for investigating molecular mechanisms of complex brain disorders. Genes Brain Behav. 13: 13–24.

Garcia-Reyero, N., B. S. Jayasinghe, K. J. Kroll, T. Sabo-Attwood, and N. D. Denslow, 2018 Estrogen signaling through both membrane and nuclear receptors in the liver of Fathead Minnow. General and Comparative Endocrinology 257: 50–66.

Gillis, J., and P. Pavlidis, 2012 “Guilt by association” is the exception rather than the rule in gene networks. PLoS Comput Biol 8: e1002444.

Horreo, J. L., 2017 Revisiting the mitogenomic phylogeny of Salmoninae: new insights thanks to recent sequencing advances. PeerJ 5: e3828–10.

Huang, D. W., B. T. Sherman, and R. A. Lempicki, 2009 Systematic and integrative analysis of large gene lists using DAVID bioinformatics resources. Nat Protoc 4: 44–57.

Kodama, M., M. S. O. Brieuc, R. H. Devlin, J. J. Hard, and K. A. Naish, 2014 Comparative mapping between Coho Salmon (Oncorhynchus kisutch) and three other salmonids suggests a role for chromosomal rearrangements in the retention of duplicated regions following a whole genome duplication event. G3 - Genes|Genomes|Genetics 4: 1717–1730.

Langfelder, P., and S. Horvath, 2012 Fast R functions for robust correlations and hierarchical clustering. J. Stat. Soft. 46: 1–17.

Langfelder, P., and S. Horvath, 2008 WGCNA: an R package for weighted correlation network analysis. BMC Bioinformatics 9: 559.

Langfelder, P., R. Luo, M. C. Oldham, and S. Horvath, 2011 Is my network module preserved and reproducible? PLoS Comput Biol 7: e1001057.

Langmead, B., and S. L. Salzberg, 2012 Fast gapped-read alignment with Bowtie 2. Nat Meth 9: 357–359.

Li, H., B. Handsaker, A. Wysoker, T. Fennell, J. Ruan et al., 2009 The Sequence Alignment/Map format and SAMtools. Bioinformatics 25: 2078–2079.

Lien, S., B. F. Koop, S. R. Sandve, J. R. Miller, M. P. Kent et al., 2016 The Atlantic Salmon genome provides insights into rediploidization. Nature 533: 200–205.

Lochmiller, R. L., and C. Deerenberg, 2000 Trade-offs in evolutionary immunology: just what is the cost of immunity? Oikos 88: 87–98.

Mackay T. F., 2001 The genetic architecture of quantitative traits. Annu. Rev. Genet. 35: 303–339.

Mackay T. F. C., Stone E. A., Ayroles J. F., 2009 The genetics of quantitative traits: challenges and prospects. Nat Rev Genet 10: 565–577.

MacManes, M. D., 2014 On the optimal trimming of high-throughput mRNA sequence data. Front. Genet. 5: 13.

Macqueen, D. J., and I. A. Johnston, 2014 A well-constrained estimate for the timing of the salmonid whole genome duplication reveals major decoupling from species diversification. Proceedings of the Royal Society B: Biological Sciences 281: 20132881–20132881.

Mähler, N., J. Wang, B. K. Terebieniec, P. K. Ingvarsson, N. R. Street et al., 2017 Gene co-expression network connectivity is an important determinant of selective constraint. PLoS Genet 13: e1006402.

Miller, K. M., I. A. Gardner, R. Vanderstichel, T. Burnley, S. Li et al., 2016 Report on the performance evaluation of the Fluidigm BioMark platform for high-throughput microbe monitoring in salmon:, 1–293 p.

Mueller, A. J., E. G. Canty-Laird, P. D. Clegg, and S. R. Tew, 2017 Cross-species gene modules emerge from a systems biology approach to osteoarthritis. NPJ Syst Biol Appl 3: 13.

Nicol, B., and Y. Guiguen, 2011 Expression profiling of wnt signaling genes during gonadal differentiation and gametogenesis in Rainbow Trout. Sex Dev 5: 318–329.

Oldham, M. C., S. Horvath, and D. H. Geschwind, 2006 Conservation and evolution of gene coexpression networks in human and chimpanzee brains. Proceedings of the National Academy of Sciences 103: 17973–17978.

Parsch, J., and H. Ellegren, 2013 The evolutionary causes and consequences of sex-biased gene expression. Nature Publishing Group 14: 83–87.

Pasquier, J., C. Cabau, T. Nguyen, E. Jouanno, D. Severac et al., 2016 Gene evolution and gene expression after whole genome duplication in fish: the PhyloFish database. BMC Genomics 17: 368.

Poley, J. D., B. J. G. Sutherland, S. R. M. Jones, B. F. Koop, and M. D. Fast, 2016 Sex-biased gene expression and sequence conservation in Atlantic and Pacific salmon lice (Lepeophtheirus salmonis). BMC Genomics 17: 483.

Prokkola, J. M., M. Nikinmaa, M. Lewis, K. Anttila, M. Kanerva et al., 2018 Cold temperature represses daily rhythms in the liver transcriptome of a stenothermal teleost under decreasing day length. J. Exp. Biol. 221: jeb170670.

Qiao, Q., S. Le Manach, B. Sotton, H. Huet, E. Duvernois-Berthet et al., 2016 Deep sexual dimorphism in adult medaka fish liver highlighted by multi-omic approach. Sci. Rep. 6: 32459.

Quinlan, A. R., and I. M. Hall, 2010 BEDTools: a flexible suite of utilities for comparing genomic features. Bioinformatics 26: 841–842.

Quinn, T. P., and C. J. Foote, 1994 The effects of body size and sexual dimorphism on the reproductive behavior of sockeye salmon, Oncorhynchus nerka. Animal Behaviour 48: 751–761.

R Core Team, 2018 R: A language and environment for statistical computing. R Foundation for Statistical Computing.

Roberts, A., and L. Pachter, 2013 Streaming fragment assignment for real-time analysis of sequencing experiments. Nat Meth 10: 71–73.

Robertson, F. M., M. K. Gundappa, F. Grammes, T. R. Hvidsten, A. K. Redmond et al., 2017 Lineage-specific rediploidization is a mechanism to explain time-lags between genome duplication and evolutionary diversification. Genome Biol. 18: 111.

Robinson, M. D., and A. Oshlack, 2010 A scaling normalization method for differential expression analysis of RNA-seq data. Genome Biol. 11: R25.

Robinson, M. D., D. J. McCarthy, and G. K. Smyth, 2010 edgeR: a Bioconductor package for differential expression analysis of digital gene expression data. Bioinformatics 26: 139–140.

Rose, N. H., F. O. Seneca, and S. R. Palumbi, 2015 Gene networks in the wild: identifying transcriptional modules that mediate coral resistance to experimental heat stress. Genome Biology and Evolution 8: 243–252.

Rowe, L., S. F. Chenoweth, and A. F. Agrawal, 2018 The Genomics of Sexual Conflict. The American Naturalist 192: 274–286.

Sauvage, C., M. Vagner, N. Derôme, C. Audet, and L. Bernatchez, 2012a Coding gene single nucleotide polymorphism mapping and quantitative trait loci detection for physiological reproductive traits in Brook Charr, Salvelinus fontinalis. G3 - Genes|Genomes|Genetics 2: 379–392.

Sauvage, C., M. Vagner, N. Derôme, C. Audet, and L. Bernatchez, 2012b Coding gene SNP mapping reveals QTL linked to growth and stress response in Brook Charr (Salvelinus fontinalis). G3 - Genes|Genomes|Genetics 2: 707–720.

Sreenivasan, R., J. Jiang, X. Wang, R. Bártfai, H. Y. Kwan et al., 2014 Gonad differentiation in Zebrafish is regulated by the canonical wnt signaling pathway. Biology of Reproduction 90: e34397–10.

Sutherland, B. J. G., T. Gosselin, E. Normandeau, M. Lamothe, N. Isabel et al., 2016 Salmonid chromosome evolution as revealed by a novel method for comparing RADseq linkage maps. Genome Biology and Evolution 8: 3600–3617.

Sutherland, B. J. G., K. C. Hanson, J. R. Jantzen, B. F. Koop, and C. T. Smith, 2014 Divergent immunity and energetic programs in the gills of migratory and resident Oncorhynchus mykiss. Mol. Ecol. 23: 1952–1964.

Sutherland, B. J. G., K. W. Koczka, M. Yasuike, S. G. Jantzen, R. Yazawa et al., 2014 Comparative transcriptomics of Atlantic Salmo salar, Chum Oncorhynchus keta and Pink Salmon O. gorbuscha during infections with salmon lice Lepeophtheirus salmonis. BMC Genomics 15: 200.

Sutherland, B. J. G., C. Rico, C. Audet, and L. Bernatchez, 2017 Sex chromosome evolution, heterochiasmy, and physiological QTL in the salmonid Brook Charr Salvelinus fontinalis. G3 - Genes|Genomes|Genetics 7: 2749–2762.

Thompson, D., A. Regev, and S. Roy, 2015 Comparative analysis of gene regulatory networks: from network reconstruction to evolution. Annu. Rev. Cell Dev. Biol. 31: 399–428.

Tritchler, D., E. Parkhomenko, and J. Beyene, 2009 Filtering Genes for Cluster and Network Analysis. BMC Bioinformatics 10: 193–9.

Vainio, S., M. Heikkilä, A. Kispert, N. Chin, and A. P. McMahon, 1999 Female development in mammals is regulated by Wnt-4 signalling. Nature 397: 405–409.

van Dam, S., U. Võsa, A. van der Graaf, L. Franke, and J. P. de Magalhães, 2017 Gene co-expression analysis for functional classification and gene-disease predictions. Brief Bioinform.

van der Most, P. J., B. de Jong, H. K. Parmentier, and S. Verhulst, 2010 Trade-off between growth and immune function: a meta-analysis of selection experiments. Functional Ecology 25: 74–80.

van Nas, A., D. Guhathakurta, S. S. Wang, N. Yehya, S. Horvath et al., 2009 Elucidating the role of gonadal hormones in sexually dimorphic gene coexpression networks. Endocrinology 150: 1235–1249.

Warnes, G. R., B. Bolker, L. Bonebakker, R. Gentleman, W. Hubert et al., 2016 gplots: Various R Programming Tools for Plotting Data.

Wijchers, P. J., and R. J. Festenstein, 2011 Epigenetic regulation of autosomal gene expression by sex chromosomes. Trends Genet. 27: 132–140.

Wong, R. Y., M. M. McLeod, and J. Godwin, 2014 Limited sex-biased neural gene expression patterns across strains in Zebrafish (Danio rerio). BMC Genomics 15: 905–10.

Wright, A. E., M. Fumagalli, C. R. Cooney, N. I. Bloch, F. G. Vieira et al., 2018 Male-biased gene expression resolves sexual conflict through the evolution of sex-specific genetic architecture. Evolution Letters 2: 52–61.

Wu, T. D., and C. K. Watanabe, 2005 GMAP: a genomic mapping and alignment program for mRNA and EST sequences. Bioinformatics 21: 1859–1875.

