## Supplemental Results for "Sex-specific co-expression networks and sex-biased gene expression in the salmonid Brook Charr *Salvelinus fontinalis*"

The following text, figures and tables support the main text.

***Methodology in developing networks and sample overviews***

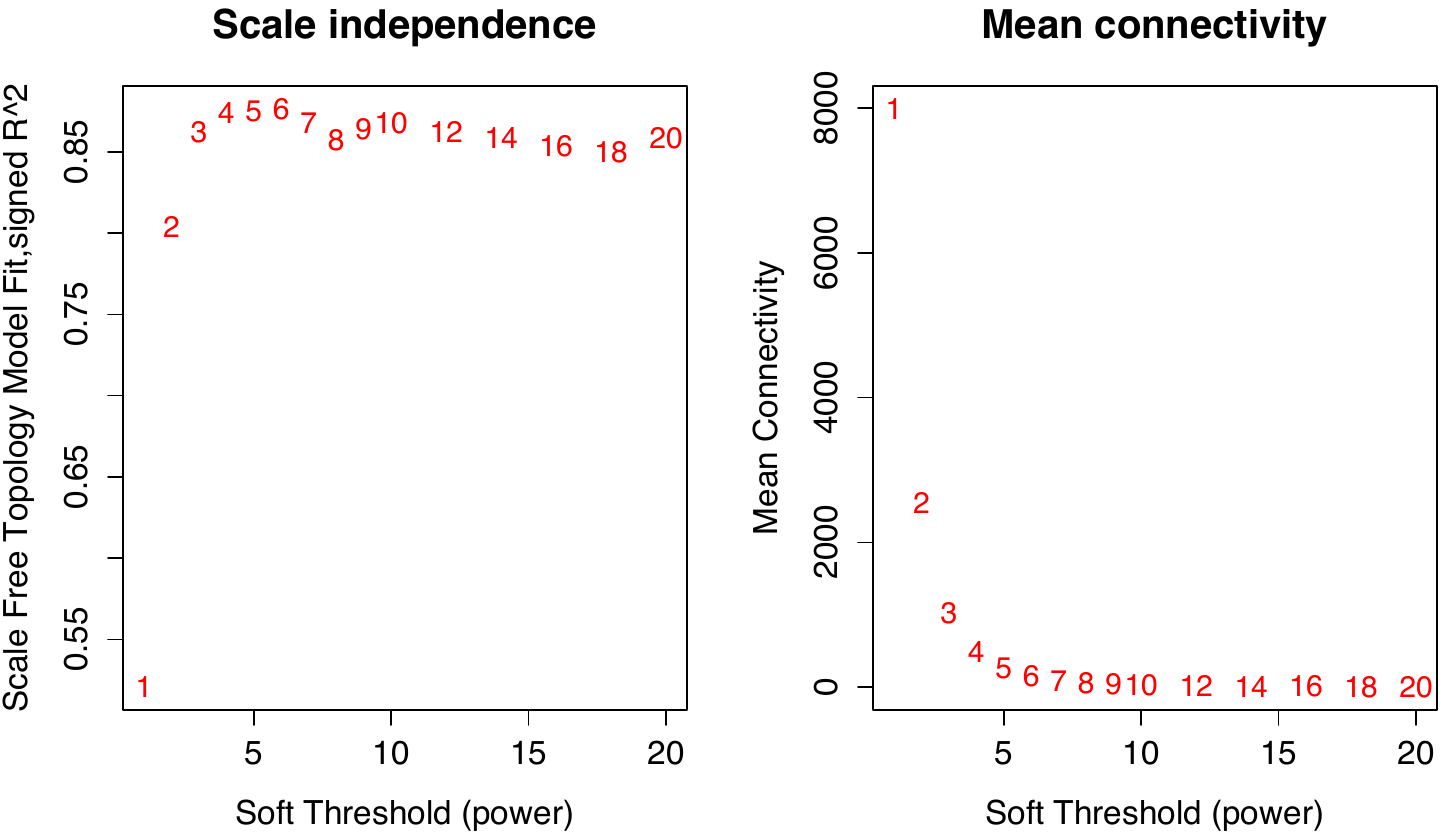

**Figure S1.** Soft threshold power selection in the female Brook Charr network after outliers were removed. Both (A) scale independence and (B) mean connectivity are used to determine the best soft threshold power. Scale free topology was chosen as beta1=6, which also is the default for the WGCNA package.

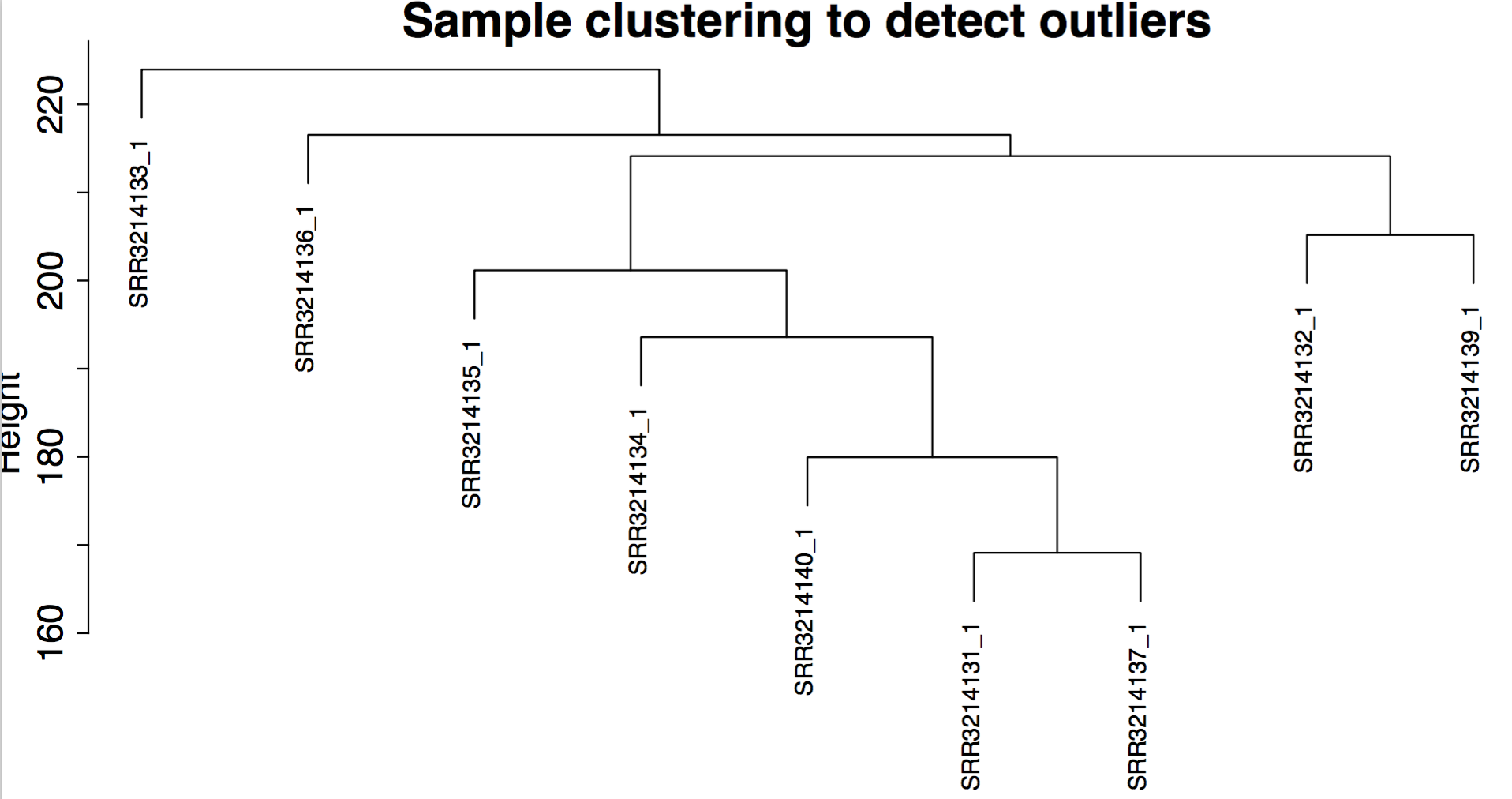

**Figure S2.** Nine individual male Arctic Charr samples clustered by gene expression similarity. Only the samples from the control condition (8 ^o^C; shown here) were used to avoid large experimental effects on the data and to make experimental conditions as parallel as possible with the Brook Charr data.

**A)**
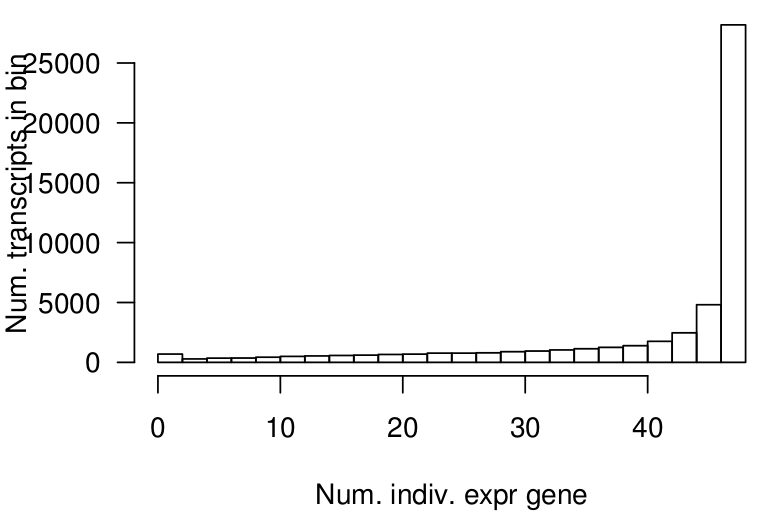
**B)**
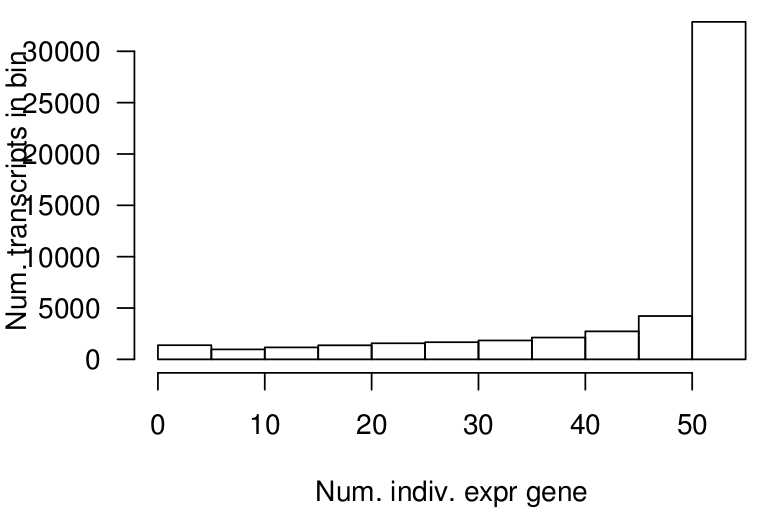

**Figure S3**. The number of transcripts expressed in the number of individuals in the Brook Charr (A) females and (B) males. This indicates that most genes were expressed in most samples within a sex, or that very few genes were specifically expressed in a small number of individuals.

**A)**

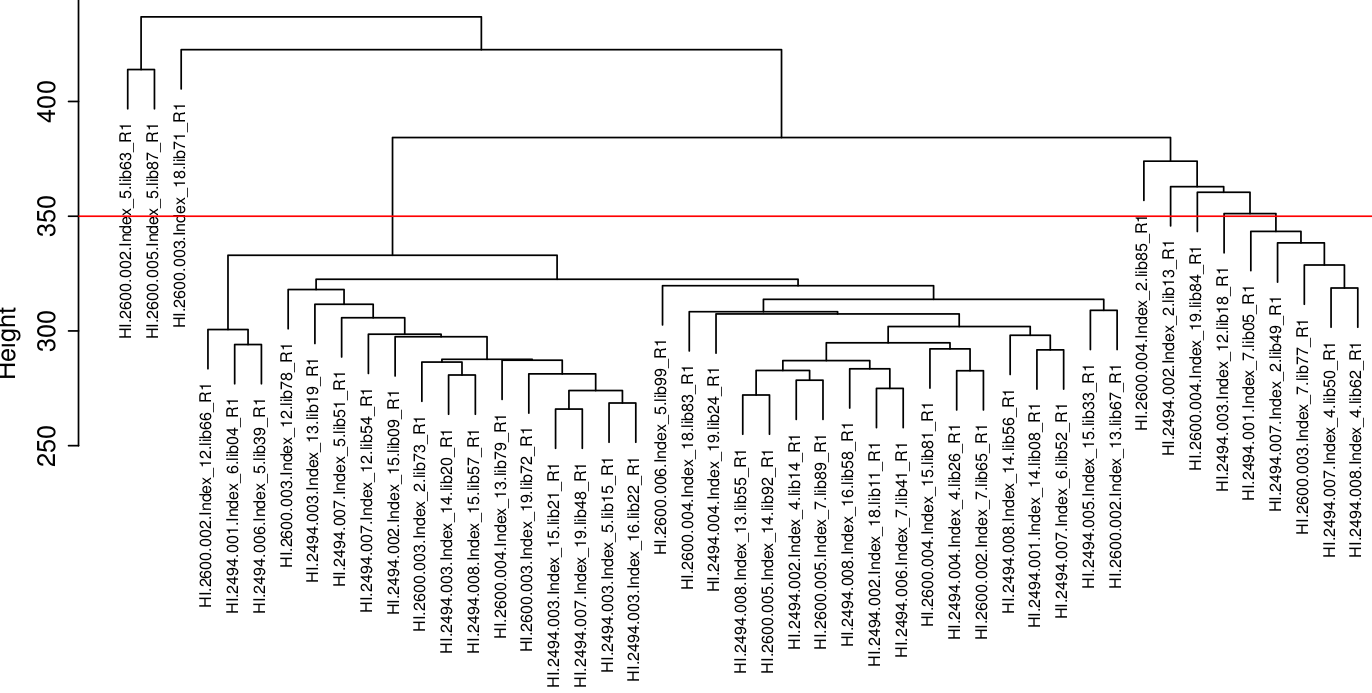

**B)**

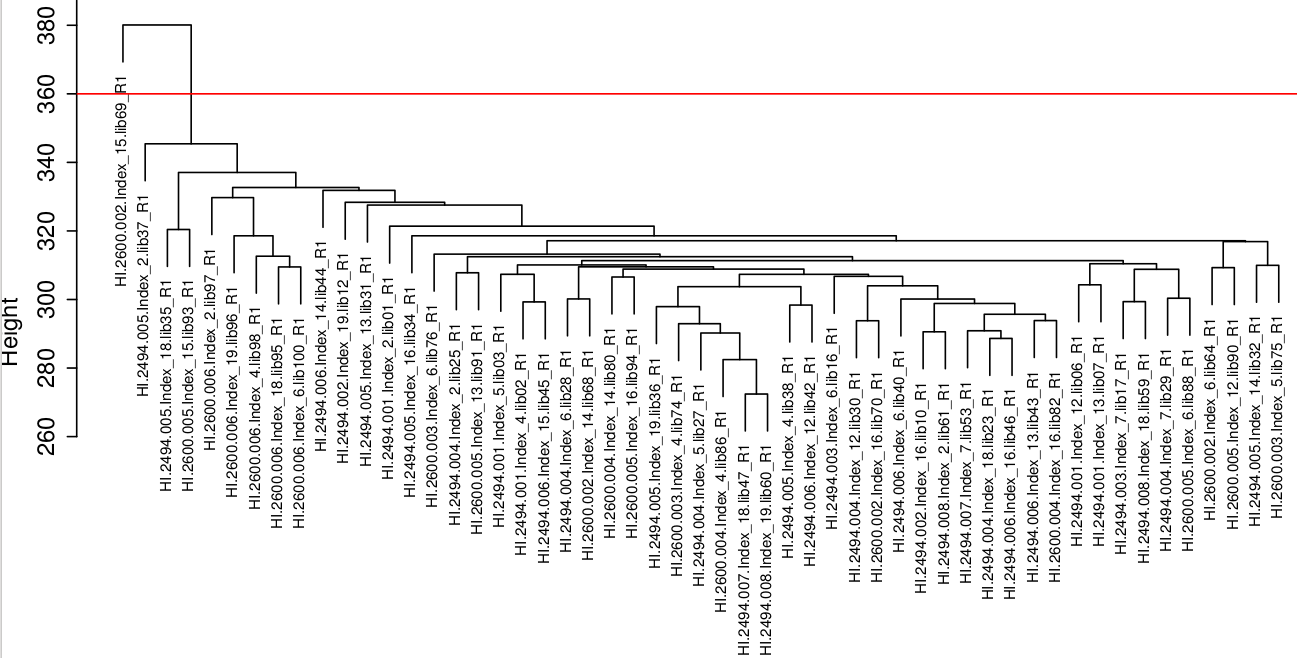

**Figure S4**. Clustering of samples by transcriptome similarity within (A) females and (B) males. Outliers were removed by discarding samples that did not group with the rest of the samples according to the cut height shown by the red horizontal line. Selection of cut height was determined by observing the clustering of samples with phenotypes and removing groups that did not fit with the rest of the samples, which included one extreme outlier male (also see Figure 1) and a subset of females with large liver weight. See Methods for further discussion on justification for outlier removal.

***Phenotypic variance differences between the sexes***

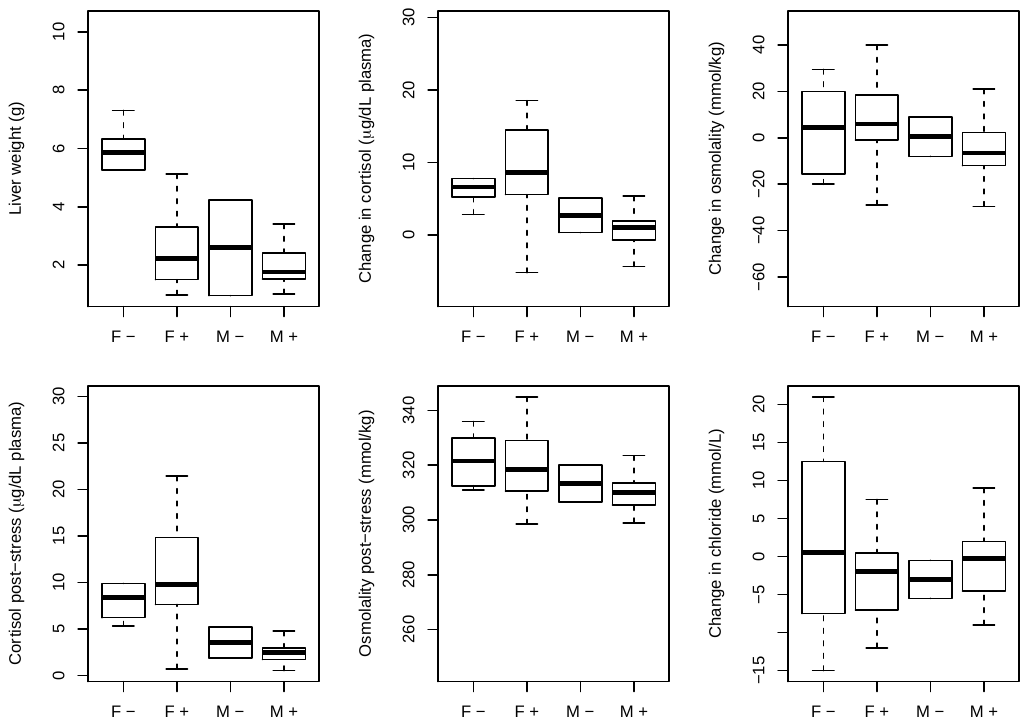

**Figure S5**. Phenotypes showed higher variance in females than males, including liver weight, osmolality and cortisol after stress, and osmolality, chloride and cortisol change from the stress exposure. Please note that there are only a few individuals in the immature category (-) relative to most individuals in the mature category (+). Outlier individuals that were removed from the network analysis are shown in blue.

***Merging modules for the female and male networks***

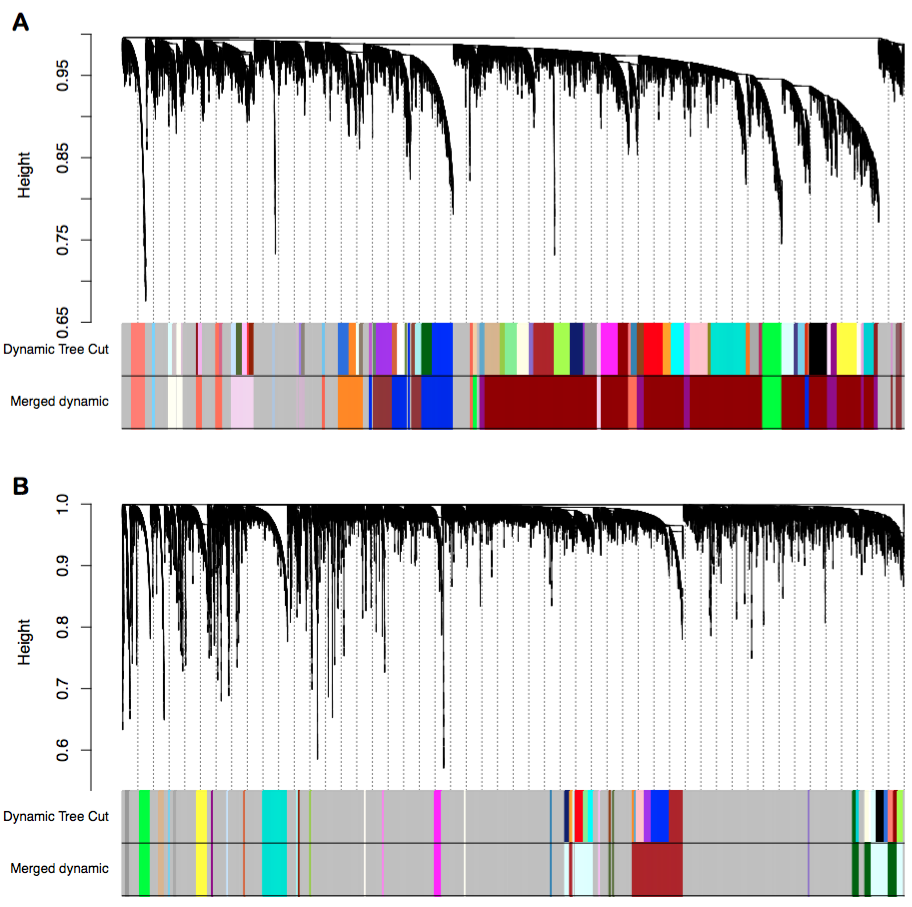

**Figure S6**. Transcripts clustered by co-expression shown as similarity to neighbouring gene (correlation between transcripts is termed ‘Height’ here) shown as vertical lines (top) and shown with assigned module (bottom) in (A) females and (B) males. Modules are shown prior to merging (Dynamic Tree Cut) and after merging similar clusters (Merged dynamic). More transcripts were assigned to modules in females (76%) than in males (28%). Note: transcript order is not retained and module colors are not related in (A) and (B).

**A)**

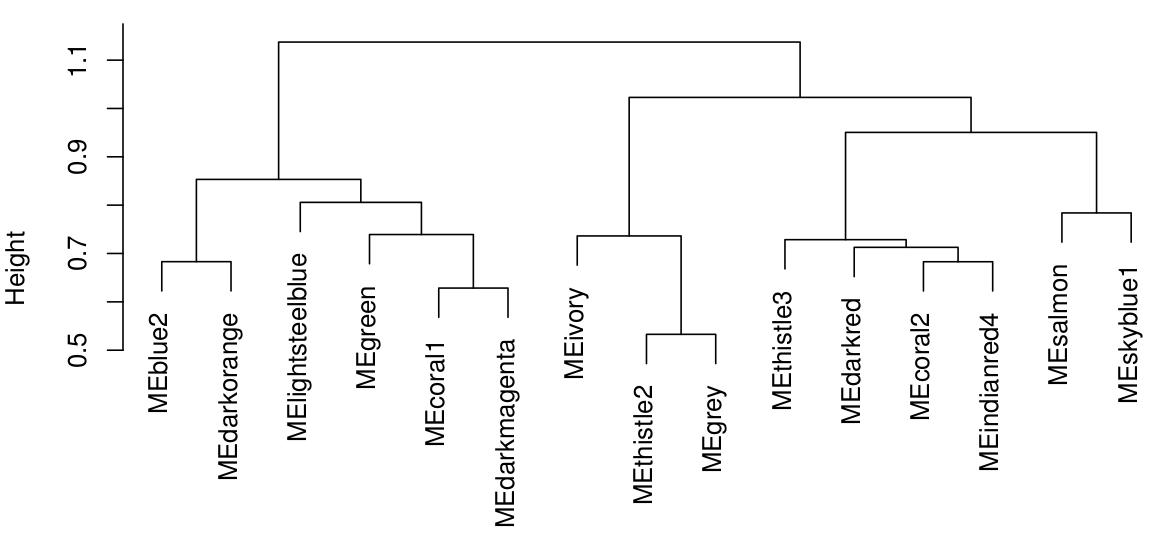

**B)**

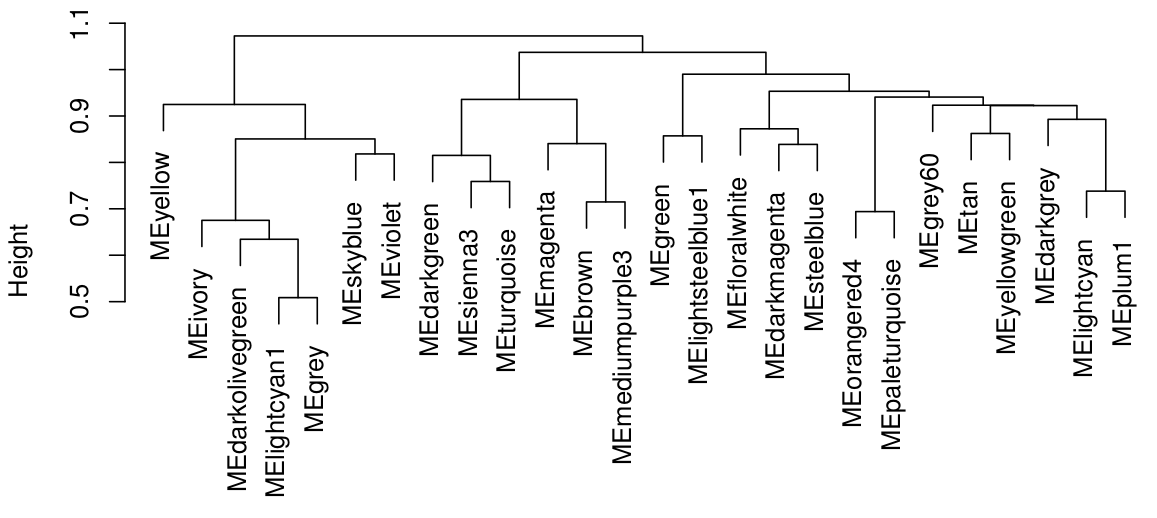

**Figure S7.** Merged modules clustered by eigengene similarity in (A) female and (B) male Brook Charr.

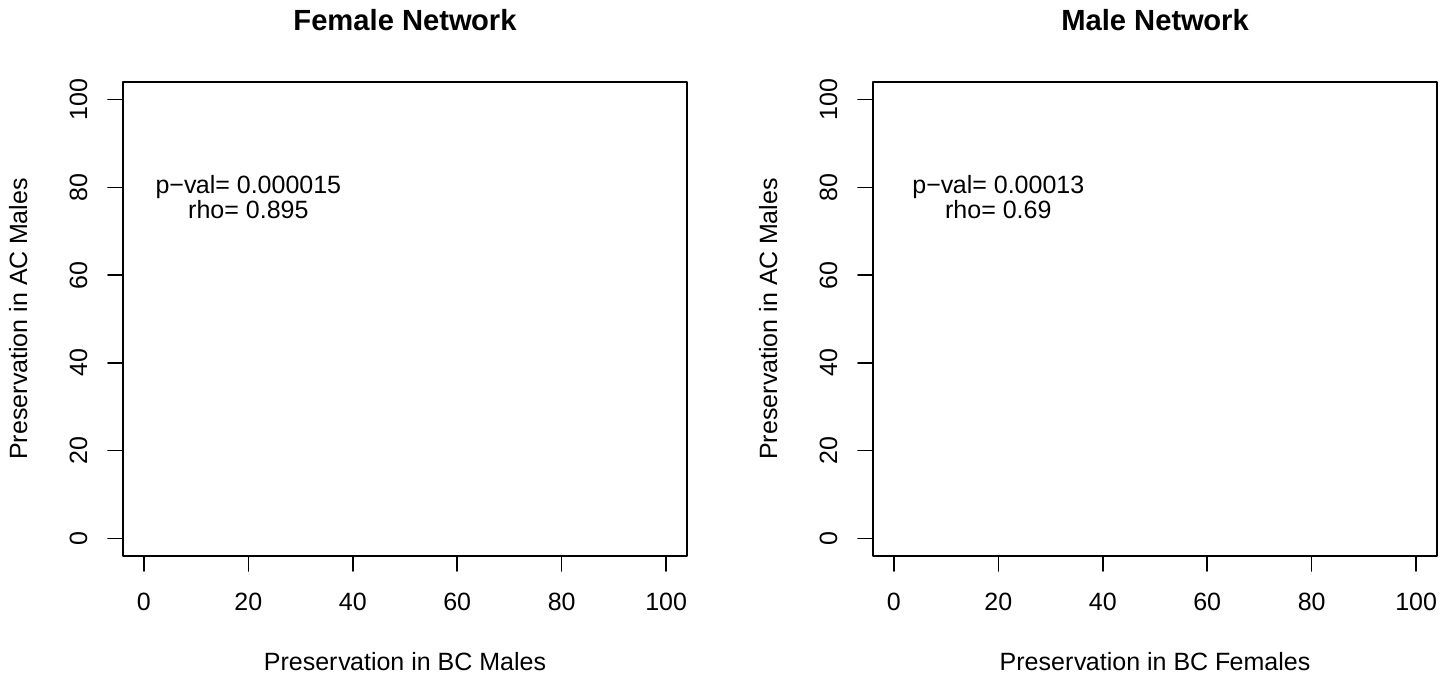

**Figure S8**. Spearman rank correlation of the module preservation statistics in either the female network (A) or the male network (B) indicating that the preservation of modules was similarly ranked in the two species.

***Cross-tabulation comparisons of female and male modules***

Cross-tabulation was used to determine the correspondence of modules and comparison of network topology between the sexes. Most importantly, 12 of the 25 male modules had more than 25% of their genes contained within the largest female module *darkred* (containing 10,533 transcripts; Additional File S2). This could be expected given the size of the *darkred* module, but this means that the genes from these male modules were often grouped together into one very large module in females. For instance, the female *darkred* module included 87% of the genes in male *darkmagenta* (virus defense response). Even though the adjacencies of the genes within male *darkmagenta* were conserved in females (preservation Zsummary = 27; Table 2), the grouping of these genes was specific to the topology of the male network. In other cases, female modules were made up of a large number of genes from a single male module. For example, the female module *lightsteelblue* (intracellular signal transduction) was comprised of a majority of genes (82%) from male *grey60*.

By contrast, 20 of the 25 assigned male modules each did not contain more than 10% of the genes from any one female module (Additional File S2). Therefore, the grouping of many male modules into the single female module *darkred* did not occur for the female network. Instead, the genes within the female modules often were largely present in the unassigned *grey* module of the males (Additional File S2). This further indicates that the male network was less assigned to modules than the female network, as these specific genes from female modules were unassigned in the male network.

***Male network module-phenotype correlation***

Similar to females, liver weight was highly correlated with modules in males, including *darkgreen* (r = 0.53) and *yellow* (r = - 0.52). Other highly correlated module-phenotype comparisons included osmolality change with *lightcyan1* (r = 0.52), *yellow* and *steelblue* (r = -0.4). Relative to the female comparisons, there were more modules strongly correlated with growth rate including *yellow* and *steelblue* (r > |-0.39|), *darkgreen* and *turquoise* (r > 0.37). Post-stress chloride was correlated with *ivory* and *lightcyan1* (r > 0.4), *steelblue* and *tan* (r > |-0.42|). Although modules from the female network did not show significant correlations with length and weight, male modules *darkgrey* and *green* were correlated with length (r > 0.42). The male-specific phenotype sperm concentration was associated with *steelblue*, *brown* (r > 0.38), and *lightcyan1* (r = -0.38).

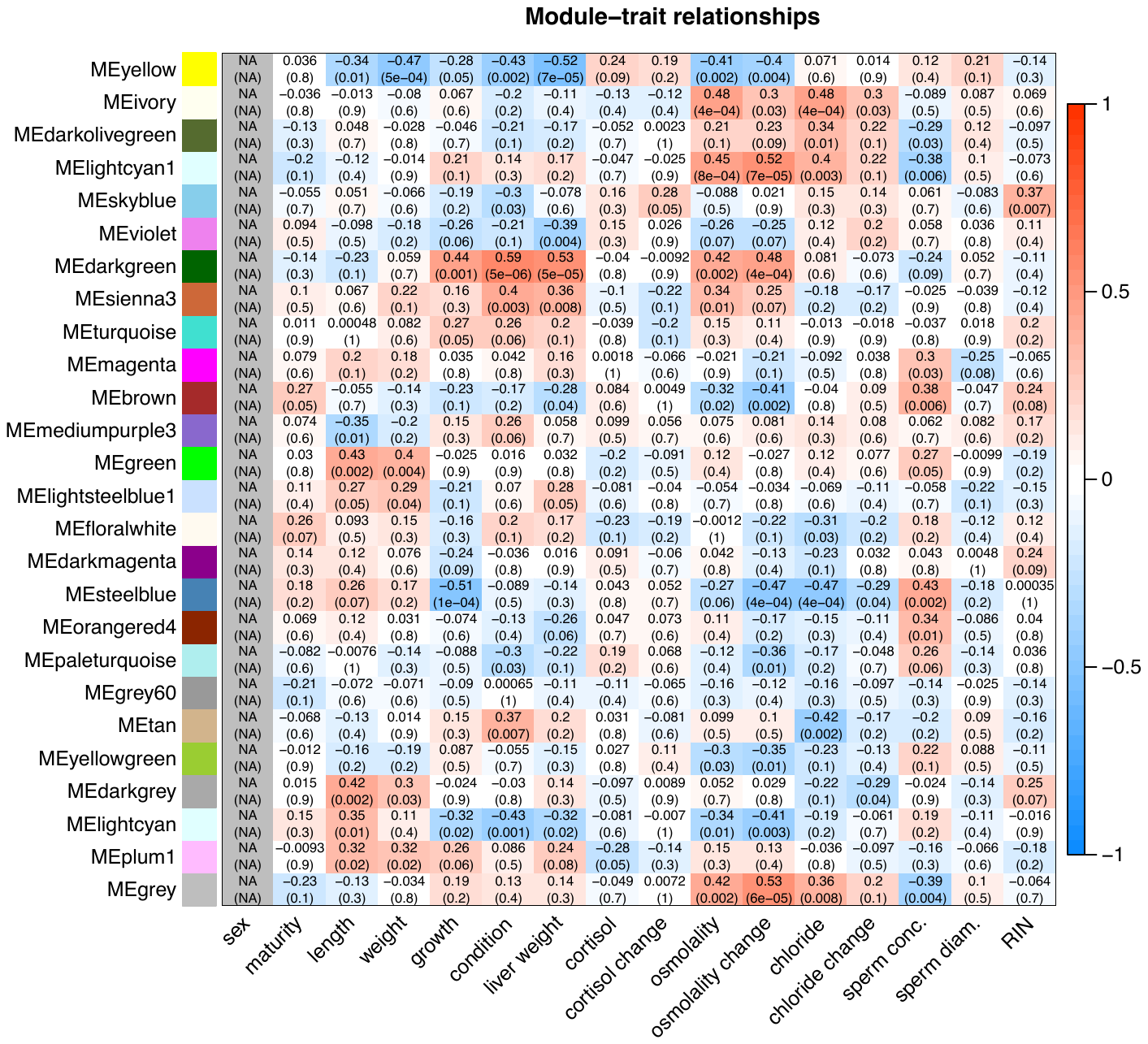

**Figure S9**. Module-trait relationships for Brook Charr males, estimated with Pearson correlation r-values and p-values. The boldness of the red or blue color indicates the strength of the relationship. Module-trait correlations are also shown in Table 2 with more general grouping of traits alongside other metrics such as module size and enriched Gene Ontology categories. Female module-trait relationships are shown in Figure 2.

***Hub genes in modules of interest***

Hub genes can be identified using module membership (MM) scores that estimate gene connectivity to other genes within the module (Additional Files S4 and S5). Importantly, the distribution of MM is continuous and there is no discrete change between a hub gene and a non-hub gene. Here we present the highest ranked hub genes for several modules from both the female and male networks that are of interest due to GO enrichment, module conservation or phenotypic correlation.

Female *lightsteelblue* was of interest due to conservation in the male network (77 transcripts; intracellular signal transduction), but hub genes (MM > 0.9) were often unannotated. The six of 15 that were annotated, included *selenium-binding protein,* several stonustoxin subunits (proteins with hemolytic activity), *estrogen-related receptor gamma*, and *NACHT, LRR and PYD domains-containing protein 12-like.* Several other transcripts, still with high MM (> 0.80), were annotated as *neoverrucotoxin subunit beta-like*, another toxin-related protein with hemolytic activity. The female module *salmon* (449 transcripts; hemopoiesis) was also of interest as it was conserved in males. Hub genes from *salmon* had 53 transcripts with MM > 0.9, most of which were annotated, often with hemopoietic functions including *band 3* (anion transport protein in erythrocytes), *erythropoietin receptor* (promotes blood cell proliferation and prevents apoptosis)*,* various hemoglobin subunits, and *5-aminolevulinate synthase* (involved in heme synthesis). A transcript annotated as the transcription regulator *Kruppel-like factor 4* also was in the top MM genes. The highest MM is *tubulin beta-6 chain-like*, which is a major constituent of microtubules.

For male modules, *ivory* was of interest due to enrichment for transcription factor activity, male-specificity of the module, and correlation with chloride levels. For this module, there were no genes with MM > 0.9. The highest MM transcripts included transcription regulators such as *frizzled-9* (FZD9) and *protein wnt* (WNT9) the ligand for the frizzled family of transmembrane receptors (MM = 0.79), the transcription factor *RAR-related orphan receptor gamma 2 protein* and *lysine-specific demethylase 4B* (*KDM4B*), which plays a role in the histone code, and transcription activator *nuclear factor 1 A-type*. Although these genes related to transcription factor activity were all ranked high in MM, there were also other transcripts lower in MM rank that are also putatively involved in transcription regulation, including those annotated as putative zinc finger proteins, or related to the SOX family of transcription factors. Male *darkmagenta* was also of interest as it was highly conserved and enriched for innate viral immunity-related functions. Only four transcripts were MM > 0.9, but some of the top ranking MM transcripts were associated to the module function, including the most connected *probable ATP-dependent RNA helicase* (DDX58/RIG-1; MM = 0.93), important for sensing viral infection and inducing type I interferons and pro-inflammatory cytokines, *interferon-induced protein with tetratricopeptide repeats 5* (IFIT5; MM = 0.89), an interferon induced RNA-binding immunity protein, *galectin-3-binding protein A* (L3BPA; MM = 0.89), *sacsin* (SACS; MM = 0.88) and *probable E3 ubiquitin-protein ligase* (HERC6; MM = 0.84) both involved in immunity in fish, and *signal transducer and activator of transcription 1-alpha/beta* (STAT1; MM = 0.83), a transcription activator that mediates responses to interferons. Male *steelblue* is of interest due to conservation and enrichment of immunity-related functions. Only one transcript had MM ≥ 0.9, *40S ribosomal protein S11-like*. However, other high-ranking hub genes included innate immunity-related genes, namely *C-type lectin domain family 4 member E* (CLC4E; MM = 0.86), *metalloreductase* (STEA4; MM = 0.86) involved in integrating inflammatory and metabolic responses. Similar to above, other lectin-related transcripts were also found lower in MM, including others annotated as *CLC4E* and as *leukocyte cell-derived chemotaxin-2* (LECT2), which has neutrophil chemotactic activity. Male *turquoise* was also of interest as it was enriched for immune system processes and had 17 hub genes, 11 of which were related to immune system processes. There were no correlated traits for this module. The interplay of these three different immune modules can be investigated by looking at module eigengene clustering (Figure S7B). The immune-related *steelblue* and *darkmagenta* modules had similarly clustering module eigengenes, and these two clustered separately from the *turquoise* module (see Figure S7B), suggesting a more distinct expression profile for *turquoise*. Male modules *yellow* (translation) and *brown* (translation) were among the most conserved across species and had correlations to phenotypes. The four hub genes of *yellow* all encode ribosomal proteins. *Brown* (related to plasma osmolality and sperm concentration) had all 22 hub genes annotated (Additional File S5). Of these hub genes, 19 were related to RNA metabolism and other metabolic processes. Two members of heat shock 70 kDa protein family members were included as hub genes in this module, *hsp4* and *hsp14*.

***Sex-specific module enrichment on the putative sex chromosome***

**Table S1. Enrichment of sex-specific modules on the putative sex chromosome.** Importantly, all genes shown below were found non-overlapping and positioned on a chromosome.

| Chromosome | Module | Fisher exact test P-value | Genes in sex chromosome in module | Genes not in sex chromosome in module |
| --- | --- | --- | --- | --- |
| NC_027308.1 | *darkgrey* | 0.59 | 1 | 13 |
| NC_027308.1 | *ivory* | 1.0 | 1 | 23 |
